# Transdifferentiation of fibroblasts into muscle cells to constitute cultured meat with tunable intramuscular fat deposition

**DOI:** 10.1101/2023.10.26.564179

**Authors:** Tongtong Ma, Ruimin Ren, Jianqi Lv, Ruipeng Yang, Zheng Xinyi, Yang Hu, Guiyu Zhu, Heng Wang

## Abstract

Current studies on cultured meat mainly focused on the muscle tissue reconstruction in vitro, but lack the formation of intramuscular fat which is a crucial factor in determining taste, texture and nutritional contents. Therefore, incorporating fat into cultured meat is of superior value. In this study, we employed the myogenic/lipogenic transdifferentiation of chicken fibroblasts in 3D to produce muscle mass and deposit fat into the same cells without the co-culture or mixture of different cells or fat substances. The immortalized chicken embryonic fibroblasts were implanted into the hydrogel scaffold and the cell proliferation and myogenic transdifferentiation were conducted in 3D to produce the whole-cut meat mimics. Compare to 2D, cells grown in 3D matrix showed elevated myogenesis and collagen production. We further induced fat deposition in the transdifferentiated muscle cells and the triglyceride content could be manipulated to match and exceed the levels of chicken meat. The gene expression analysis indicated that both lineage-specific and multi-functional signalings could contribute to the generation of muscle/fat matrix. Overall, we were able to precisely modulate muscle, fat, and extracellular matrix contents according to balanced or specialized meat preferences. These findings provide new avenues for customized cultured meat production with desired intramuscular fat contents that can be tailored to meet the diverse demands of consumers.

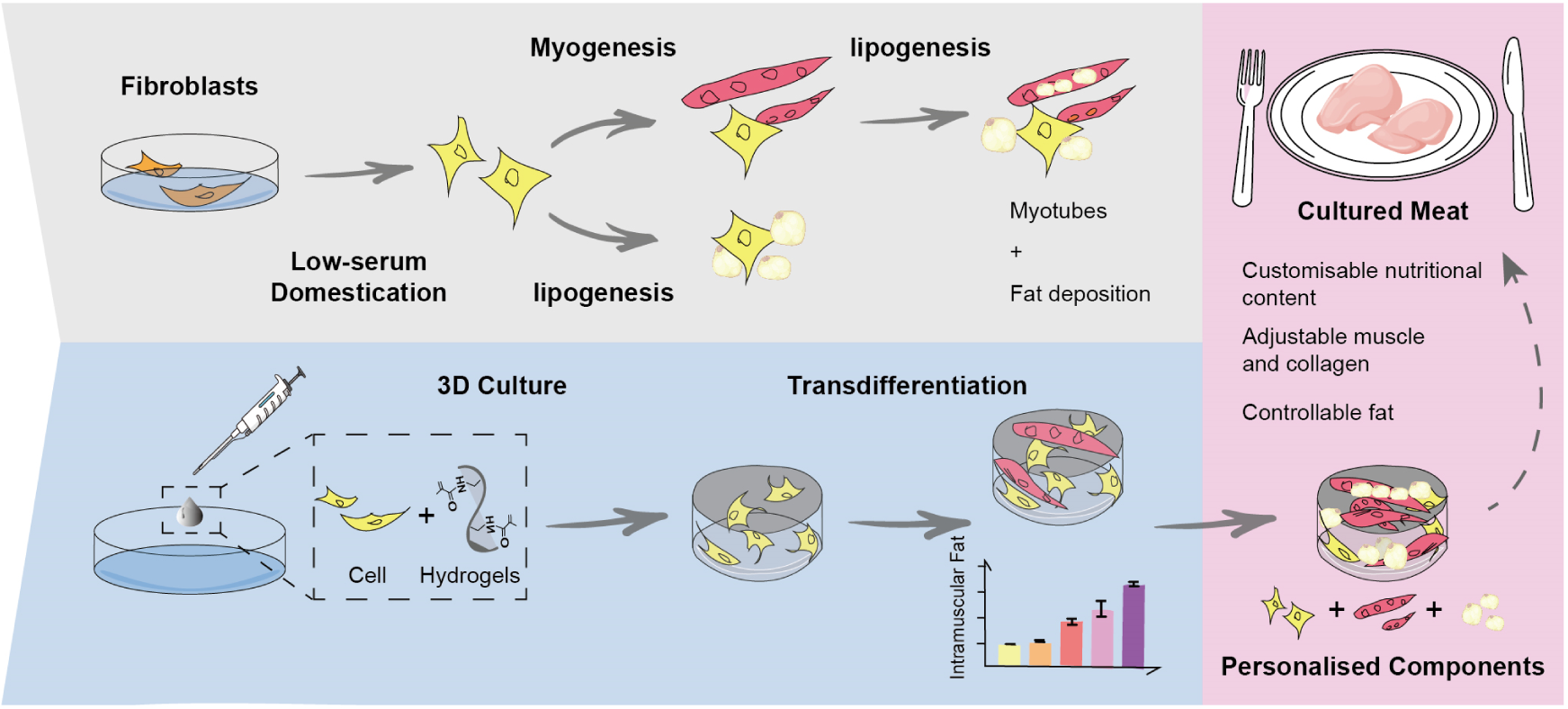
Graphical Abstract.

## Introduction

Cultured meat is an innovative and emerging technique that produces meat directly from cell cultures, potentially providing a high-quality, safe, and stable source of animal protein (Chriki and Hocquette, 2020). In comparison to traditional livestock and poultry farming, cultured meat generates fewer greenhouse gases, utilizes less arable land, and causes less animal harm (Mattick et al., 2015). Proper selection of seed cells is one of the keys to the success of cultured meat production. The starting cells must be easily obtainable and be able to proliferate numerous times to enable mass production. While the stem cells like muscle stem cells and pluripotent stem cells have frequently been utilized as the cellular source for cultured meat, these stem cells are rare in the animal body and difficult to obtain and amplify on a large scale. In contrast, the somatic cells, which constitute the body, could be efficiently converted into muscle cells under certain conditions. The fibroblast is one of the most abundant and widely-distributed cell types present in the body and could be easily collected via minimally invasive biopsy procedure without sacrifice the farm animals. The fibroblasts can replicate indefinitely in vitro and are amenable for myogenesis, adipogenesis and chondrogenesis (French et al., 2004; Yin et al., 2010), which produce muscle, fat and extracellular matrix (ECM) proteins that constitute the meat and the associated texture and flavor. Recently, the fibroblast cells from farm animals have been utilized as the source cells for cultured meat production, with or without myogenesis (Jeong et al., 2022; Pasitka et al., 2023), demonstrating the feasibility of somatic cell-derived seed cells as a sustainable and ethical option for cultured meat production.

We have previously developed a protocol for the controlled transdifferentiation of chicken fibroblasts into myoblasts, which subsequently form multinucleated myotubes and express mature muscle proteins (Ren et al., 2022). Moreover, chicken fibroblasts could also be efficiently converted into fat-depositing lipocytes by treating them with chicken serum medium or other substances such as fatty acids (Hausman, 2012; Kim et al., 2020; Lee et al., 2021). Therefore, the induced myogenic and adipogenic competency, along with the inherent fibrogenic collagen-producing ability of chicken fibroblasts, can enable us to simultaneously synthesize muscle, fat and collagen during the cultured meat production. Nevertheless, further techno-functional research is necessary to precisely control the composition of the end product of fibroblast-derived cultured meat in order to achieve a more balanced and personalized nutritional profile and meet the specific consumer preferences.

In this study, the chicken fibroblast cells were implanted into the hydrogel scaffold to analyze the 3D cellular dynamics involving cell proliferation and myogenic/adipogenic transdifferentiation. We also optimized the low-serum culture conditions of chicken fibroblasts to reduce the cost of mass production. The myogenic transdifferentiated cells were confirmed to be skeletal muscle lineage but not myofibroblasts. Importantly, the cells were subjected to myogenesis and adipogenesis sequentially in 3D hydrogel matrix to resemble the whole-cut meat with the controllable intramuscular fat and collagen content. By using transdifferentiation strategies, the depositing ratio of fat into the cultured meat could be manually and precisely adjusted, allowing the nutrients to be naturally synthesized from the organized muscle structure. This demonstrates the potential for manipulating cultured meat to meet consumer preferences for specific fat content and texture.

## Results

### Chicken fibroblasts proliferate stably in low-serum conditions

The chicken fibroblast cells were chosen as the ideal cell source for cultured meat production because they can propagate indefinitely and undergo myogenesis whenever the induction is provided. These cells can be readily obtained from fertilized eggs without the need to harvest animals. We have previously constructed an inducible myogenic transdifferentiation system in chicken fibroblasts with the stable integration of Tet-On-MyoD cassette (Ren et al., 2022). The MyoD is the key myogenic transcription factor and the chicken fibroblasts could be converted into striated and elongated myotubes (myofibers) upon forced expression of MyoD (Ren et al., 2022; Weintraub et al., 1989). The Tet-On-MyoD cassette enables the inducible and reversible activation of chicken MyoD factor, and in the current fibroblast cells, the myogenic transdifferentiation is only switched on by adding doxycycline (DOX). Without the DOX treatment, the control chicken fibroblast cells (Tet-On-MyoD) do not differ from the wild-type cells in terms of morphology, proliferation rate and gene expression (Figure S1A, S1B, S1C). Hence, as a proof-of-concept experiment, we utilized this inducible myogenic fibroblast cell line to develop protocols for cultured chicken meat production.

To achieve sustainable and cost-effective cell production, it is essential to minimize the serum usage in the culture medium. Therefore, we implemented a progressive serum reduction approach to acclimate the cells in the prospect of obtaining a stable cell source that can be propagated in low serum concentrations. In the initial experiment, we used 12% fetal bovine serum (FBS) in 1640 basal medium, which served as the control group. The results showed that the cells were able to proliferate normally at 6%, 3% and 1.5% serum concentrations, but at decreasing rates as shown by the EdU assay (Figure 1A and 1B). We also tested the chicken serum as an alternative to bovine serum, in order to avoid cross-species contamination of animal derivatives. The results showed that chicken fibroblasts could proliferate stably in the 1% chicken serum (Figure 1A, 1B, S1D) and the cell populations multiply 3 times in 3 days as demonstrated by the CCK-8 assay (Figure 1C). Thus, we conclude that the low-serum medium can effectively support the stable propagation of chicken fibroblasts.

**Figure 1.**
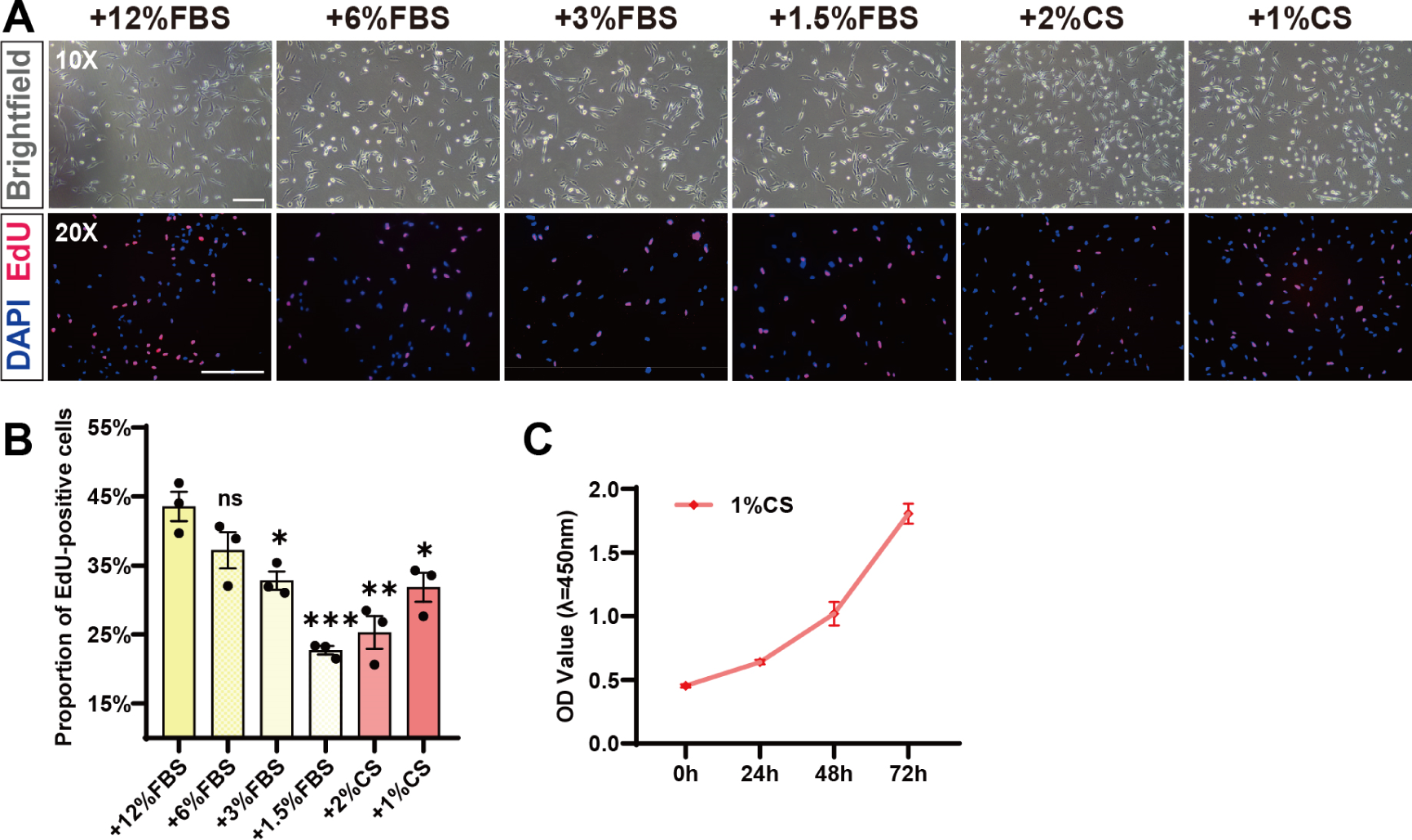
Chicken fibroblasts proliferate stably in low-serum conditions. (A) Cellular morphology and EdU staining of chicken fibroblasts under different low-serum conditions. FBS: Fetal bovine serum. CS: Chicken serum. Scale bar, 200 µm. (B) Quantification of the proportion of EdU-positive cells in Figure A. Error bars indicate s.e.m. n = 3. *P < 0.05, **P < 0.01, ***P < 0.001. (C) The CCK-8 cell proliferation assay showed the proliferation of chicken fibroblasts in 1% chicken serum. Error bars indicate s.e.m, n = 3.

### 3D culture of chicken fibroblasts in GelMA hydrogels

The behavior of cells grown on the top of 2D flat surface may differ from that of cells in 3D space. To simulate the 3D natural growth environment of cells *in vivo*, we utilized the Gelatin Methacrylate (GelMA)-based hydrogels as scaffolds for chicken fibroblasts. GelMA hydrogel can form a stable and porous structure for cell implantation and is commonly used in tissue engineering because of its great biocompatibility and mechanical tenability (Pepelanova et al., 2018). We created hydrogels with varying concentrations at 3%, 5%, 7%, 9%, and then observed their surface characteristics when immersed in culture medium using an emission scanning electron microscope. The porosity was measured to be 83.27%, 65.01%, 62.47% and 57.96%, respectively. Thus, the higher the mass concentration of hydrogels, the tighter the 3D mesh structure formed and the smaller the porosity (Figure 2A). The pore diameters in the scaffolds were determined to range from 3µm^2^ to 100µm^2^, showing the biggest pore size in 3% hydrogel and smallest in 9% hydrogel (Figure 2B). Therefore, the porosity and pore size of the scaffold could be adjusted to achieve optimal physical strength and nutrients delivery that suited for long-term cell culturation.

**Figure 2.**
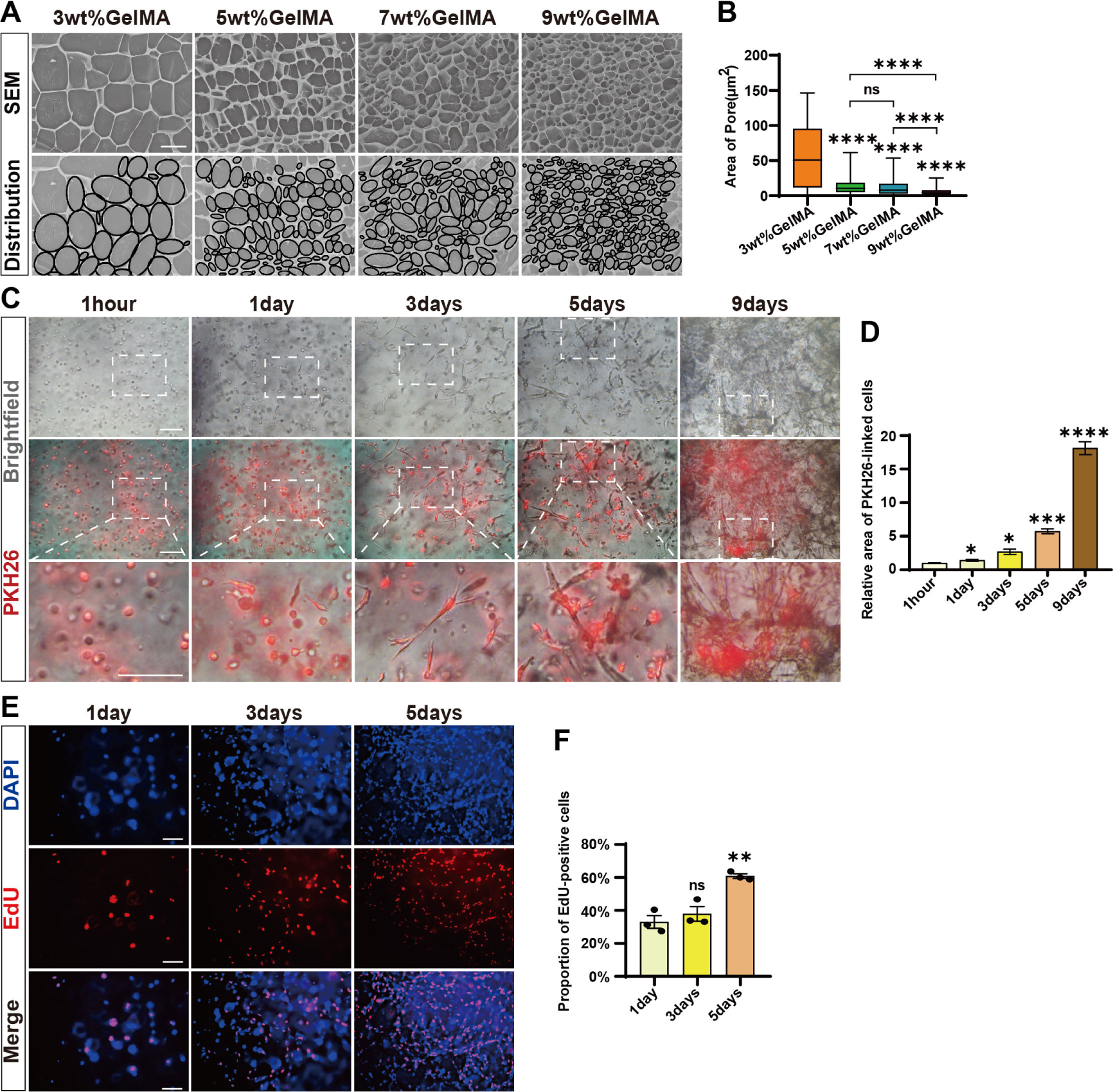
3D culture of chicken fibroblasts in GelMA hydrogels. (A) Microscopic images of GelMA hydrogels at different concentrations (3%wt, 5%wt, 7%wt and 9%wt) taken by scanning electron microscopy and their corresponding simplified maps of pore distributions. Scale bar, 10 µm. (B) Quantification of pore area in image A. Error bars indicate s.e.m, n = 3. ****P < 0.0001. (C) Brightfield and red fluorescent images of cells in 3D culture after PKH26 staining at different times (1hour, 1day, 3days, 5days and 9days). Scale bar, 100 µm. (D) Relative area of PKH26-linked cells in Figure C. Error bars indicate s.e.m, n = 3. *P < 0.05, ***P < 0.001, ****P < 0.0001. (E) Representative EdU staining shows the proliferation of cells in 3D culture on 1day, 3days and 5 days after cell implantation in hydrogel. Scale bar, 100 µm. (F) Quantification of the proportion of EdU-positive cells in Figure E. Error bars indicate s.e.m, n = 3. **P < 0.01.

To determine the most ideal gel concentration and pore size suited for chicken fibroblast attachment and growth, we conducted an experiment where we implanted the same number of cells into gels with varied pore sizes and examined the cellular dynamics over a 9-day period (Figure S2). The cells adhered to the gels immediately and started to exhibit typical extended and irregular fibroblast cell morphology after 1 day, indicating the hydrogel’s good biological compatibility with chicken fibroblast cells. However, the 3% hydrogel collapsed in the medium 2 days after cell implantation and thus did not support the long-term cell growth. We continued to monitor the 3D cell proliferation and found that the cells successfully propagated in 5%, 7%, 9% hydrogels for the entire duration of the experiment. Among the tested hydrogels, the 5% concentration was found to be the best option for cell attachment and growth as shown by the densest cellular structure formed, and thus be utilized for the subsequent analysis.

Due to the challenges of observing and distinguishing the cells and pores within the hydrogel scaffold using the brightfield of ordinary light microscopy, we then used the PKH26 fluorescent cellular dye to accurately observe the cellular morphology and quantity the cell numbers. The cells were firstly labelled with PKH26 in 2D culture (Figure S3A) and then transferred into the gel matrix. Upon implantation, the cells appeared mostly round on the first day, but gradually extended and expanded within the hydrogel scaffold. By the 5th day, most of the cells were elongated with irregular shape and short tentacles and packed tightly, and by the 9th day they multiplied more than 15 times and formed dense fibrous bundles (Figure 2C and 2D). We also examined the proliferation dynamics of the cells in 3D scaffold via the EdU assay. The proliferation efficiency of chicken fibroblast cells gradually increased overtime in 3D culture and reached replication levels even higher than those in 2D culture (Figure 2E and 2F). This result indicates that 3D culture conditions provide a more favorable environment for cell growth. Moreover, when the 3D cultured cells were detached from the gel by collagenase dissociation (Figure S3B) and seeded back in 2D monolayer culture plates, the cells again exhibited similar morphology but slightly increased proliferation capacity compared to the original fibroblasts in 2D conditions (Figure S3C and S3D). Thus, the chicken fibroblast cells cultured in 3D maintain their normal physiological characteristics and we continue to stimulate the cellular myogenesis and lipogenesis to prepare them for cultured meat production.

### Transdifferentiation of chicken fibroblasts into muscle cells in 3D

Despite the complexity in the composition of fresh meat and processed products, muscle cells are the major and indispensable component of meat foods. The muscle cell derived myotubes (myofibers) provide a rich source of proteins and nutrients and constitute the meat texture. To this end, we aimed to transform the chicken fibroblast into muscle cells through the established MyoD overexpression protocol (Ren et al., 2022). We first tested and optimized the myogenic transdifferentiation parameters in 2D culture through the DOX induced MyoD expression (Figure S4). We observed elongated multinucleated myotubes and abundant expression of myosin heavy chain (MHC) after 3 days of myogenic induction (Figure S5A). The “serum deprivation” protocol was the classical strategy to stimulate terminal myogenic differentiation in myoblasts of various species including chicken (Nakashima et al., 2005). Thus, we examined the myogenic transdifferentiation efficiency (myotube fusion index) in the chicken fibroblast cells overexpressing MyoD in combination with reduced concentrations of bovine or horse serums. However, we found that reducing serum concentrations did not increase the myotube formation but instead caused massive cell death instead (Figure S5B). Therefore, it seems that the “serum deprivation” could not further improve the myogenic transdifferentiation in chicken fibroblast cells and a simple MyoD overexpression strategy is sufficient for efficient production of mature muscle cells.

After demonstrating the feasibility of induced transdifferentiation of chicken fibroblasts into muscle cells in 2D culture, we continued myogenic transdifferentiation in 3D to simulate the construction of concrete whole-cut meat. The cells were inoculated into a hydrogel scaffold and allowed to proliferate for 7 days before inducing transdifferentiation with MyoD overexpression (Figure 3A). We used whole-amount MHC immunofluorescence staining to examine the myotube formation directly inside the gel. We identified abundant multinucleate MHC^+^ myotubes at multiple focal planes within the gel (Figure 3B, Supplementary Video 1), indicating successful myogenic transdifferentiation of 3D cultured chicken fibroblasts. In contrast, there were no MHC signal or autofluorescence detected in the 3D cultured chicken fibroblasts (Figure S5C) and hydrogel without cells (Figure S5D). With the assistance of confocal Z-stack analysis, the stacked images showed densely packed MHC+ myotubes from a piece of cellular hydrogel complex at a depth of 68 μm (Figure 3D). The separate XY axis views of the orthogonal projections at different depths (Figure 3E) and a multi-angle video (Supplementary Video 2) also showed the several myotubes were aligned together. Nevertheless, many myotubes were oriented in different directions, preventing the entire matrix from aligning in one direction. In conclusion, we successfully transformed the chicken fibroblast cells into mature muscle cells in 3D environment.

**Figure 3.**
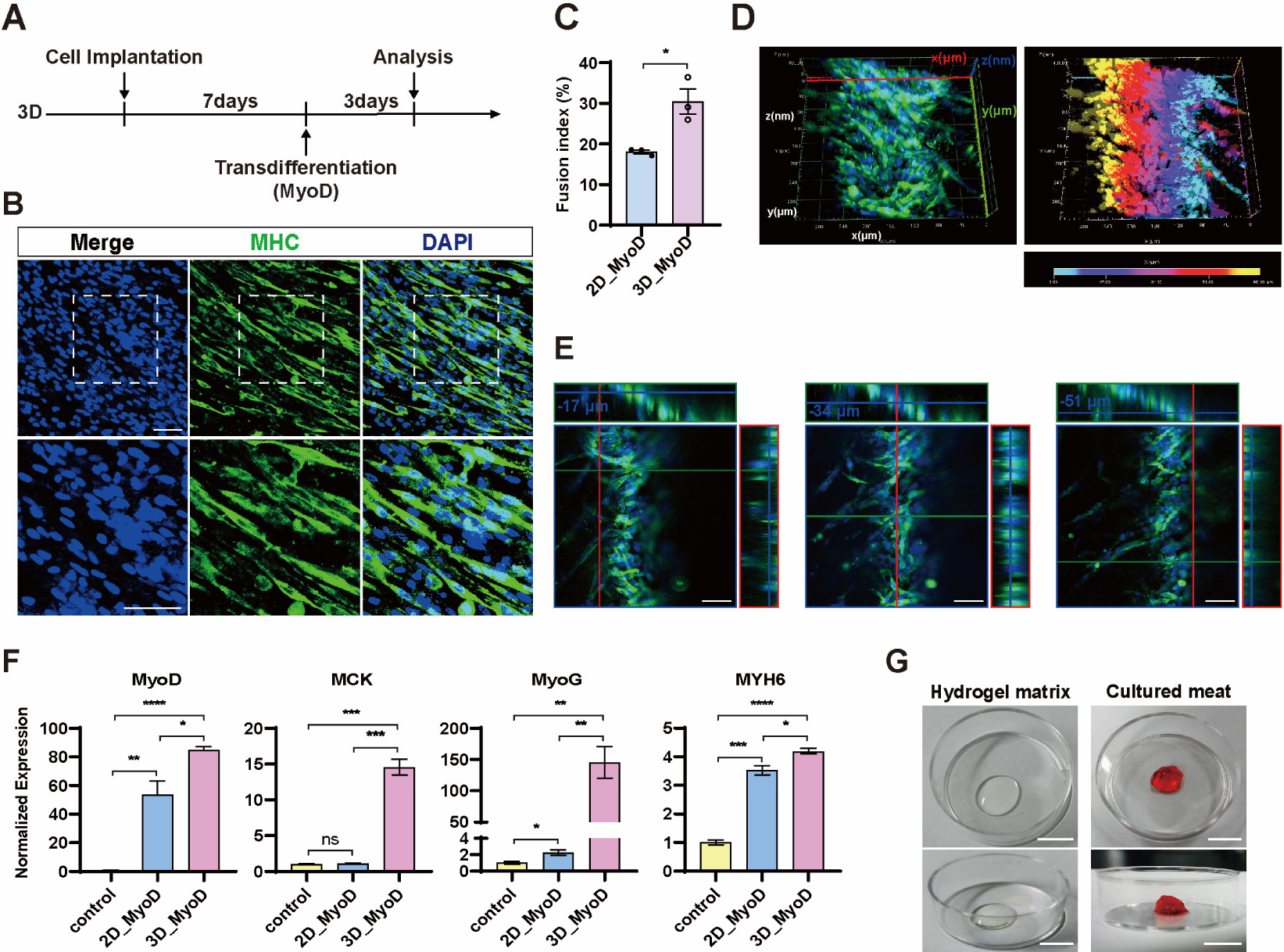
Transdifferentiation of chicken fibroblasts into muscle cells in 3D. (A) Experimental design for fibroblast myogenic transdifferentiation in 3D culture. (B) Representative images of MHC staining showed the myogenic ability of chicken fibroblasts in 3D culture. Scale bar, 50 µm. (C) Comparison of the mean myogenic fusion index between 2D and 3D. Error bars indicate s.e.m, n = 3. *P < 0.05. (D) Three-dimensional images of MHC staining of cells cultured in 3D. The right panel is its depth-coded image which indicate different depths from the deepest (cyan) to the surface (yellow). (E) Orthogonal projections of three sets of MHC staining of cells in 3D culture at different depths. Scale bar, 50 µm. (F) Expression of skeletal muscle-related genes was determined by RT-qPCR in 2D and 3D cells upon myogenic transdifferentiation and control 3D cells without stimulation. Please note that the myogenic transdifferentiation driven by MyoD stimulate the expression of classical myogenic factors. Error bars indicate s.e.m, n = 3. *P < 0.05, **P < 0.01, ***P < 0.001, ****P < 0.0001. (G) Macroscopic morphology of the empty hydrogel matrix (left) and cultured meat (right). The cultured meat is the product obtained after 3D cell culture and induction of myogenesis/lipogenesis. Scale bar, 1 cm.

Compared to 2D, the transdifferentiated myotubes induced in 3D were more organized and densely packed, resembling the native myofiber distribution *in vivo* (Figure 3B). And the myotube formation efficiency (fusion index) in 3D reached 30.49%, which was significantly higher than that of the 2D under the same transdifferentiation conditions (Figure 3C). We also evaluated the expression of several myogenic markers by RT-qPCR (Figure 3F). MyoG (Myogenin) is a transcription factor regulating terminal skeletal muscle differentiation and could be induced by MyoD (Cao et al., 2006). MYH6 (Myosin Heavy Chain 6) is the major protein comprising the muscle thick filament, and functions in muscle contraction (van Rooij et al., 2009; Warkman et al., 2012). MCK (Muscle Creatine Kinase) is an enzyme that primarily active in mature skeletal and heart muscle (Vincent et al., 1993). All myogenic factors, including MyoG, MYH6, and MCK, significantly increased upon transdifferentiation in both 2D and 3D. In addition, the transdifferentiated cells exhibited significant higher expression of MyoG and MCK in 3D conditions compared to that in 2D, indicating more robust myogenic differentiation and maturation of cells in the 3D microenvironment. We speculate that the porous structure of the hydrogel matrix may support the cells to grow in all directions, similar to the environment in which the myofibers form in vivo.

Macroscopically, the muscle filaments and the dense cellular network structures formed by myogenic transdifferentiation could make the hydrogel matrix more compact and solid (Figure 3G). Thus, compared to the empty transparent scaffolds without cells, the hydrogels cultivated with muscle cells are more visually like whole-cut meat thus resemble to the fresh animal meat.

### Myogenic transdifferentiation of fibroblast does not produce myofibroblasts

Fibroblasts can be induced to differentiate into myofibroblast upon injury, leading to tissue fibrosis. Similar to the skeletal muscle cells, the myofibroblast are also contractile and express the myogenic factor MyoD as well as certain types of sarcomeric myosin heavy chains (MHC) (Hecker et al., 2011). To confirm that the myogenic transdifferentiated cells were indeed skeletal muscle cells but not the myofibroblasts, we utilized a panel of cell lineage-specific markers to delineate the cell conversion progress and determine the cellular identity of fibroblasts, myoblasts and myofibroblasts, respectively. The immunofluorescence staining of 3D cultured cells showed that the muscle-specific intermediate filament Desmin (Paulin and Li, 2004) was expressed only in the MyoD transdifferentiated cells, but not in the original fibroblasts or the embryonic skin derived myofibroblasts (Figure 4A). In contrast, the classical myofibroblast marker alpha-smooth muscle actin (α-SMA) (Hinz et al., 2007; Takase et al., 1988) was expressed only in myofibroblasts, but not in the fibroblast or the transdifferentiated cells (Figure 4B). The fibroblast intermediate filament Vimentin (Tarbit et al., 2019) was abundantly expressed in the fibroblasts but reduced in the myogenic transdifferentiated cells (Figure 4C). The 2D and 3D cultured cells showed consistent pattern of marker protein expression, indicating that the different culture models and conditions do not affect the cell identity conversion (Figure S6A and S6B). These results confirmed that the MyoD indeed transdifferentiate the cells toward the skeletal muscle lineage but not the myofibroblast. Furthermore, the RT-qPCR showed that, after myogenic induction in 3D, the skeletal muscle specific genes Desmin and Six1 (Relaix et al., 2013) were significantly elevated (Figure 4D) whereas the fibroblast gene Thy-1 was significantly reduced (Figure 4E). The transforming growth factor β (TGFβ) signaling is the most potent known inducer of myofibroblast differentiation (Vaughan et al., 2000). We found the expression of core TGFβ signaling components, TGFβ-1, TGFβ-3 and Smad3, remain unchanged during the transdifferentiation process (Figure 4F), indicating that the classical myofibroblast lineage was not induced. Together, these data confirm that the myogenic transdifferentiation of fibroblast does not produce myofibroblasts.

**Figure 4.**
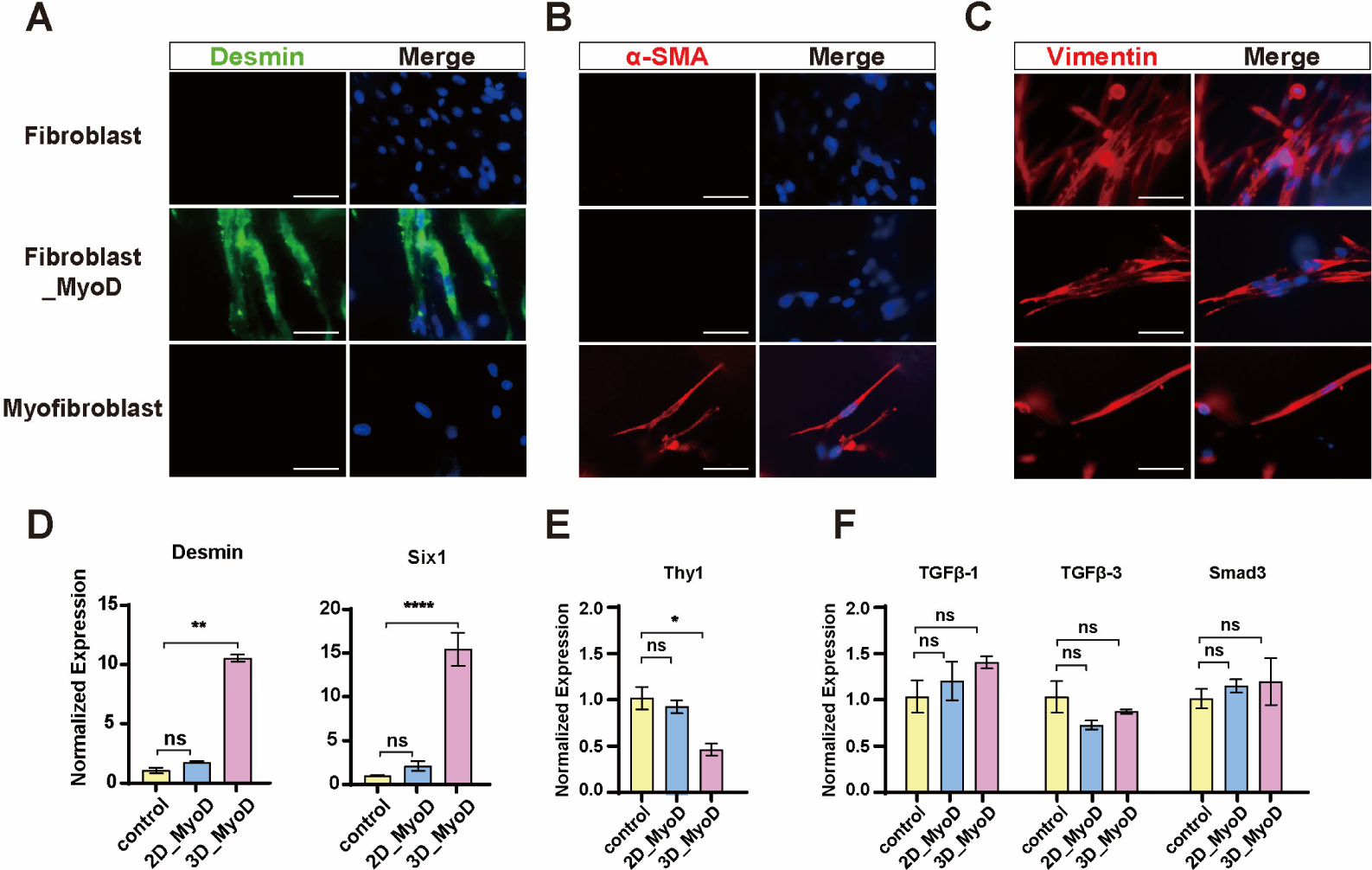
Myogenic transdifferentiation of fibroblasts do not produce myofibroblasts. (A) Immunofluorescence staining of 3D cultured cells showed that the skeletal muscle marker desmin was expressed only in the transdifferentiated cells but not in fibroblasts or myofibroblasts. Scale bar, 50 µm. (B) Immunofluorescence staining of 3D cultured cells showed that the myofibroblast marker α-SMA was expressed only in the myofibroblasts but not in fibroblasts or transdifferentiated cells. Scale bar, 50 µm. (C) Immunofluorescence staining of 3D cultured cells showed that the fibroblast marker vimentin was abundantly expressed in fibroblasts and myofibroblasts but greatly reduced in transdifferentiated cells. Scale bar, 50 µm. (D) RT-qPCR showed that the myogenic genes Desmin and Six1 were significantly increased upon myogenic transdifferentiation. (E) RT-qPCR showed the fibroblast marker gene Thy-1 was significantly reduced upon myogenic transdifferentiation. (F) The myofibroblast marker genes TGFβ-1, TGFβ-3 and Smad3 remain unchanged during myogenic transdifferentiation. Error bars indicate s.e.m, n = 4. *P < 0.05, **P < 0.01, ***P < 0.001, ****P < 0.0001. ns: not significant.

### Stimulate the fat deposition in chicken fibroblasts in 3D

The intramuscular fat is a crucial component of meat that can determine its quality attributes, such as taste and flavor. The chicken fibroblasts have been reported to be amenable for lipogenesis through various stimulus, including chicken serum, insulin, fatty acids and retinoic acids (Kim et al., 2021; Kim et al., 2020; Lee et al., 2021). We first attempted different lipogenic stimulations on 2D cultured cells and stained them for Oil Red O to visualize and quantify fat deposition in the fibroblasts. Only very few scattered Oil Red O signals were found in the treatments consisting of only serums (12.5% FBS, 1% chicken serum or 2% chicken serum), indicating no lipogenesis during normal proliferation conditions. However, when we added insulin and fatty acids (oleic/linoleic acid) to the medium, lipid droplets in the cells dramatically increased as detected by the Oil Red O staining. After extensive optimization of the concentrations of supplements in the medium, we identified that the 8µg/ml fatty acids plus 60µg/ml insulin (abbreviated as FI, F: fatty acids, I: insulin) can induce lipogenesis most efficiently in 2D chicken fibroblasts (Figure S7A, S7B and S7C).

Next, the same lipogenesis induction strategy was applied to the 3D cultured cells (Figure 5A). We found extensive Oil Red O signal inside the hydrogel matrix at different focal planes and the lipid droplets were clearly visible as beaded strings under magnification (5B and 5C, Supplementary Video 3). The RT-qPCR analysis illustrated the expression of genes involved in lipogenesis and triglyceride synthesis were significantly higher in the cells with lipogenic stimulation compare to the control (Figure 5D). Notably, the genes encoding for PPARγ (peroxisome proliferator-activated receptor gamma), Gpd1 (glycerol-3-phosphate dehydrogenase) and FABP4 (fatty acid binding protein 4) all showed higher expression in 3D than that in 2D. Interestingly, the expression of Znf423 (zinc finger protein 423), which is a PPARγ transcriptional activator (Addison et al., 2014; Longo et al., 2018), only increased upon lipogenic induction in 3D but not in 2D conditions. It seems that the cells grown in 3D hydrogel showed enhanced lipogenesis compare to the flat surface cultured cells, similar to the myogenic transdifferentiation process in 3D. In addition to the lipogenic induction with fatty acids and insulin (FI), we also validated the use of chicken serum in promoting lipid accumulation as previous reported (Kim et al., 2021). Our results showed that the chicken serum alone could also stimulate fat accumulation effectively (Figure S7D, S7E and S7F). We further measured the triglyceride content and found that the lipogenic induction increased the triglyceride significantly in the cell matrix (Figure 5E). In conclusion, the 3D cultured chicken fibroblast can efficiently deposit lipid by different stimulus.

**Figure 5.**
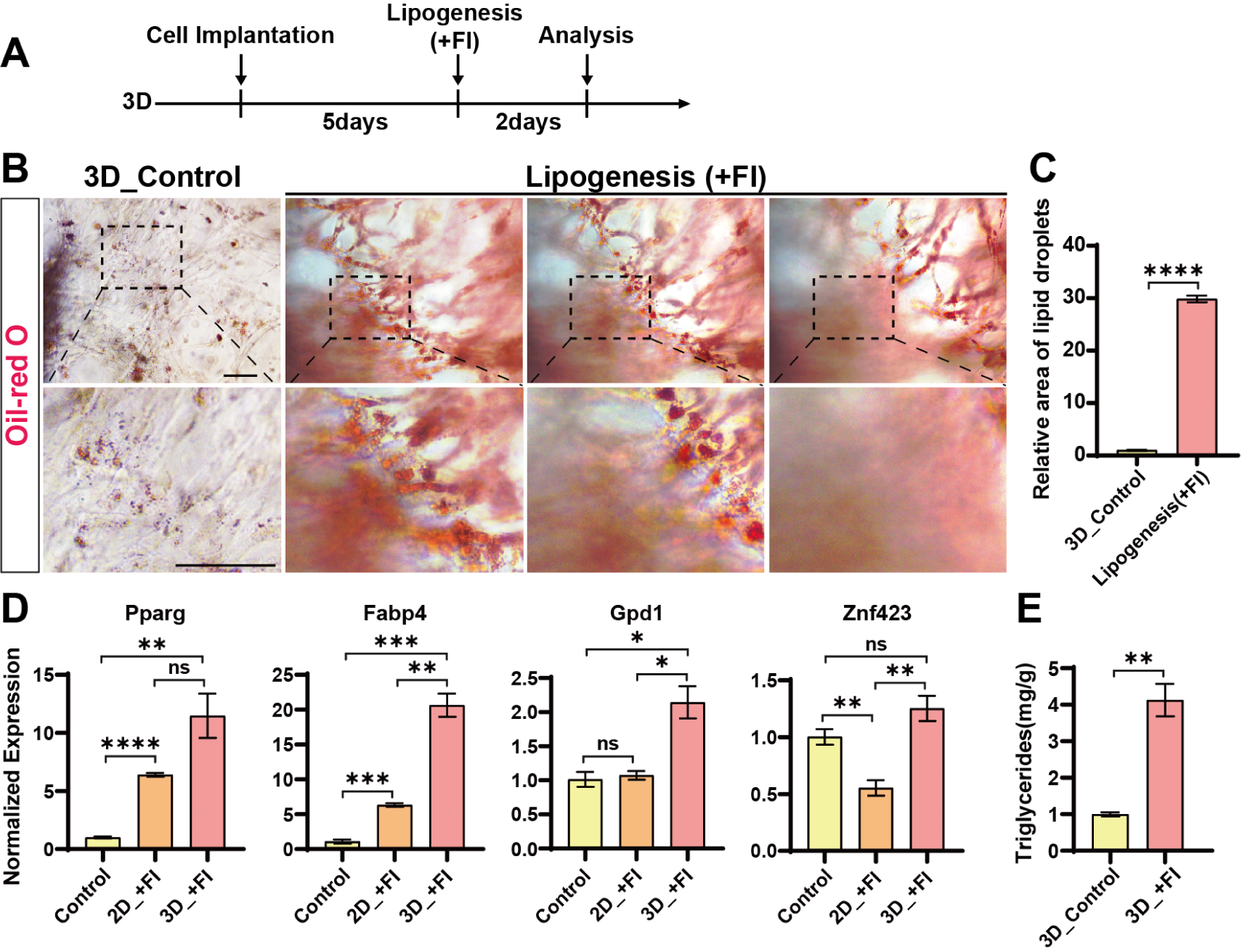
Stimulate the fat deposition in chicken fibroblasts in 3D. (A) Experimental design for fibroblast lipogenesis in 3D culture (‘F’ is for fatty acids and ‘I’ is for insulin). (B) Representative images showing the Oil-red O staining of lipid content accumulated in cells at different focal planes at the same position. The control group was normal medium without lipogenesis. Scale bar, 100 µm. (C) Relative area of lipid droplets in figure B. Error bars indicate s.e.m, n = 3. ****P < 0.0001. (D) Expression of lipid synthesis related genes determined by RT-qPCR in 2D and 3D cells upon lipogenic induction and control 3D cells without stimulation. Error bars indicate s.e.m, n = 3. *P < 0.05, **P < 0.01, ***P < 0.001, ****P < 0.0001. (E) Triglyceride content in the cultured meat upon different lipogenic inductions and control 3D cells without stimulation. Error bars indicate s.e.m, n = 3. *P < 0.05, **P < 0.01, ***P < 0.001.

### Controlled fat deposition in the transdifferentiated muscle cells in 3D hydrogel

The above presented data shows that chicken fibroblast cells have a superior capacity for transforming into muscle and depositing fat when cultured in a 3D hydrogel matrix. Next, we tried to combine the myogenic and lipogenic stimuli together to modulate the fat deposition in the cultured meat to simulate the various intramuscular fat contents in the conventionally raised meat. Rather than converting fibroblasts into muscle cells and fat cells separately and mixing them later, we adopted a new strategy that can induce de novo lipid deposition in the muscle by first inducing myogenic transdifferentiation and then followed by lipid induction in the same cells (Figure 6A and S8A). In 2D conditions, plenty of MHC^+^ myotubes and Oil Red O-stained lipids were found to intermingle after the myogenic/lipogenic treatment (Figure S8B) and some of the red marked lipid droplets were located inside the myotubes, indicating that the transformed muscle cells indeed deposit fat autonomously to constitute intramyocellular lipids (Figure S8C). Next, we applied the similar treatment to the cells cultured in 3D hydrogel and also identified Oil Red O-labelled lipid droplets mixed with the densely packed MHC^+^ myotubes (Figure 6B and 6C). These findings suggest that the use of myogenic/lipogenic treatments can induce the formation of muscle cells that are capable of depositing fat in both 2D and 3D. We further examined the expression levels of both myogenic and lipogenic factors in the 3D cultured cells by RT-qPCR. Compare to the control 3D cells without myogenic/lipogenic stimulations, the induced cells showed significantly higher expression of genes involved in both myogenesis and lipogenesis (Figure 6D). Interestingly, the extent of gene upregulation upon the combined myogenic/adipogenic stimulations was comparable to that of myogenic or adipogenic induction alone. This finding suggests that sequential myogenic transdifferentiation and lipid deposition do not interfere with each other when conducted in the same cells.

**Figure 6.**
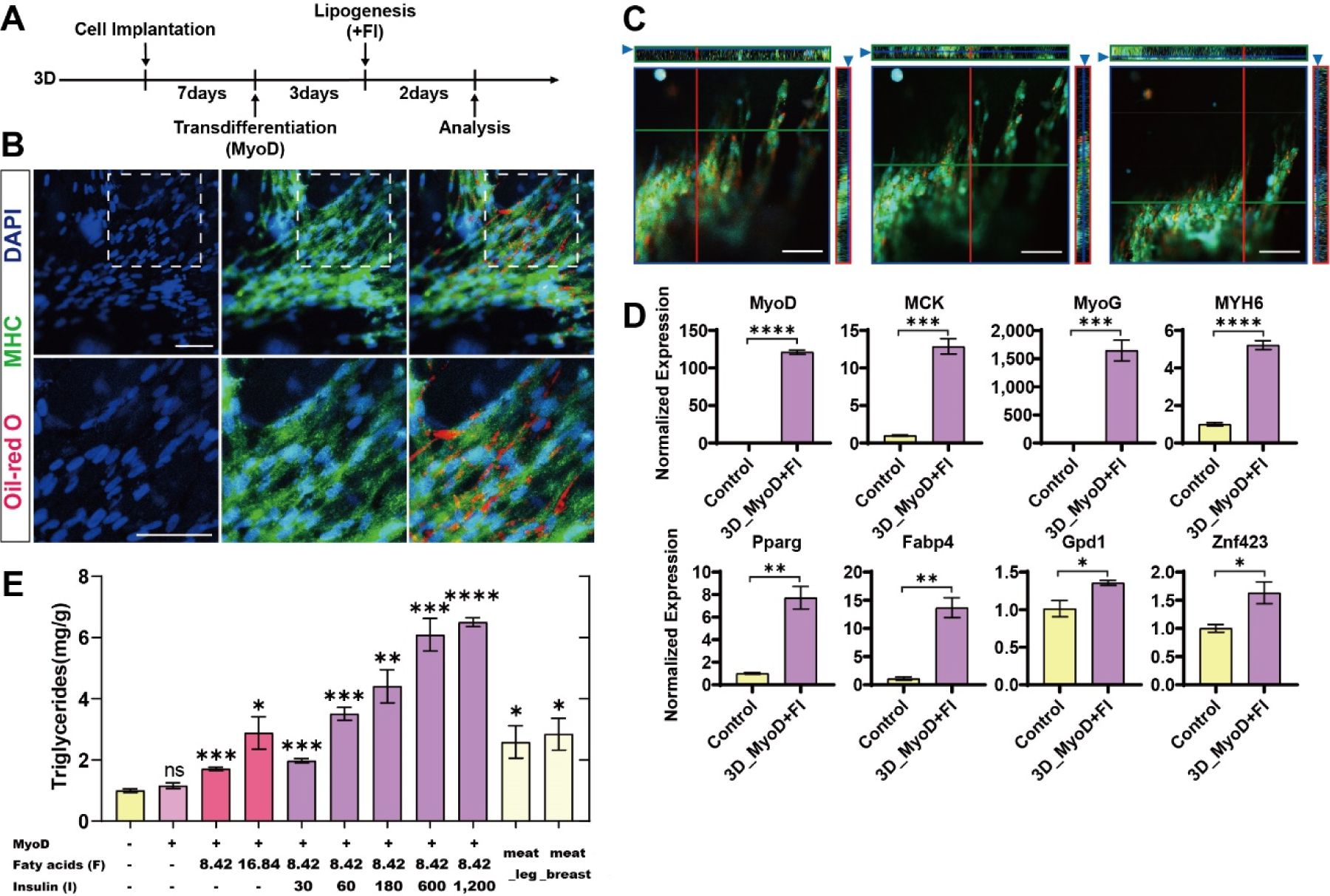
Controlled fat deposition in the transdifferentiated muscle cells in 3D hydrogel. (A) Experimental design for fibroblast myogenic/lipogenic differentiation in 3D culture. (B) Representative images of MHC and Oil Red O staining of cells upon myogenesis/lipogenesis in 3D culture. Scale bar, 50 µm. (C) Orthogonal projections of three sets of MHC and Oil Red O staining of cells in 3D culture at different depths. Scale bar, 50 µm. (D) Expression of muscle-related genes (top) and lipid-related genes (bottom) in the cells with myogenesis/lipogenesis induction and control 3D cells without any stimulation were determined by RT-qPCR. (E) Triglyceride content of cultured meat under different conditions and real meat. “Meat_leg” and “Meat_breast” are taken from the leg and breast muscles of adult chickens. Error bars indicate s.e.m, n = 3. *P < 0.05, **P < 0.01, ***P < 0.001, ****P < 0.0001.

The intramuscular fat is an integral component of both the traditional animal meat and the cultured meat, and it directly influences the meat flavor and texture (Frank et al., 2016). Hence, we compared the triglyceride levels in the 3D hydrogel cells with different types of lipogenic stimuli with the chicken breast and leg meat. The results showed that the lipogenic stimulation in the 3D hydrogel cells increased the triglyceride content in the cultured meat to the levels comparable or even higher than real chicken meat (Figure 6E). In contrast, the control cells without any induction or with only myogenic stimulation do not show apparent triglyceride accumulation (Figure 6E). Therefore, the fat content in the cultured meat could be synthesized by a controlled manner and then we tried to purposely manipulate the triglyceride contents in the meat matrix by adjusting the potency of adipogenic stimulation. By fine-tuning the concentrations of insulin and fatty acids during lipogenic induction, the triglyceride contents in the final product of cultured meat can precisely reach any customized levels across the range from 1.5mg/g to 7mg/g (Figure 6E), which overlap and surpass the levels in the fresh chicken breast and leg meat. As a result, this strategy greatly expands the diversity and category of cultured meat products, allowing for precise control over intramuscular fat contents to meet consumer preferences. Therefore, guided and graded fat deposition in cultured meat allows for the creation of various meat products with controlled intramuscular fat contents.

### The collagen content and ECM components of cultured meat

Fibroblasts are an essential source of extracellular matrix (ECM) including the collagen, which providing elasticity to the tissue in the body and enriching the texture of the cultured meat (Ben-Arye and Levenberg, 2019). In theory, the fibroblast should produce abundant ECM to produce a more realistic meat product. We then examined the collagen content in the cultured meat and found that the total collagen protein gradually increased and reached the plateau at 1.59 µg/mg in the final product (Figure 7A). This is mainly due to the increased cell numbers and the accumulation of secreted collagen in the hydrogel matrix. Nevertheless, the RT-qPCR showed that the genes encoding the major components of ECM exhibited various expression patterns with the extension of culture time. The expression of COL1A1 (Collagen, type I, alpha 1) and COL1A2 (Collagen, type I, alpha 2) gradually decreased, whereas the fibronectin increased during the time course of meat synthesis. The expression of elastin and laminin genes remain stable throughout the whole course of experiment (Figure 7B). However, the laminin protein content was accumulated and increased steadily during 3D culturation (Figure 7C). Overall, the synthesis and accumulation of different types and amounts of ECM components during the myogenic/lipogenic stimulations can improve the texture of the cultured meat prepared from fibroblast cells.

**Figure 7.**
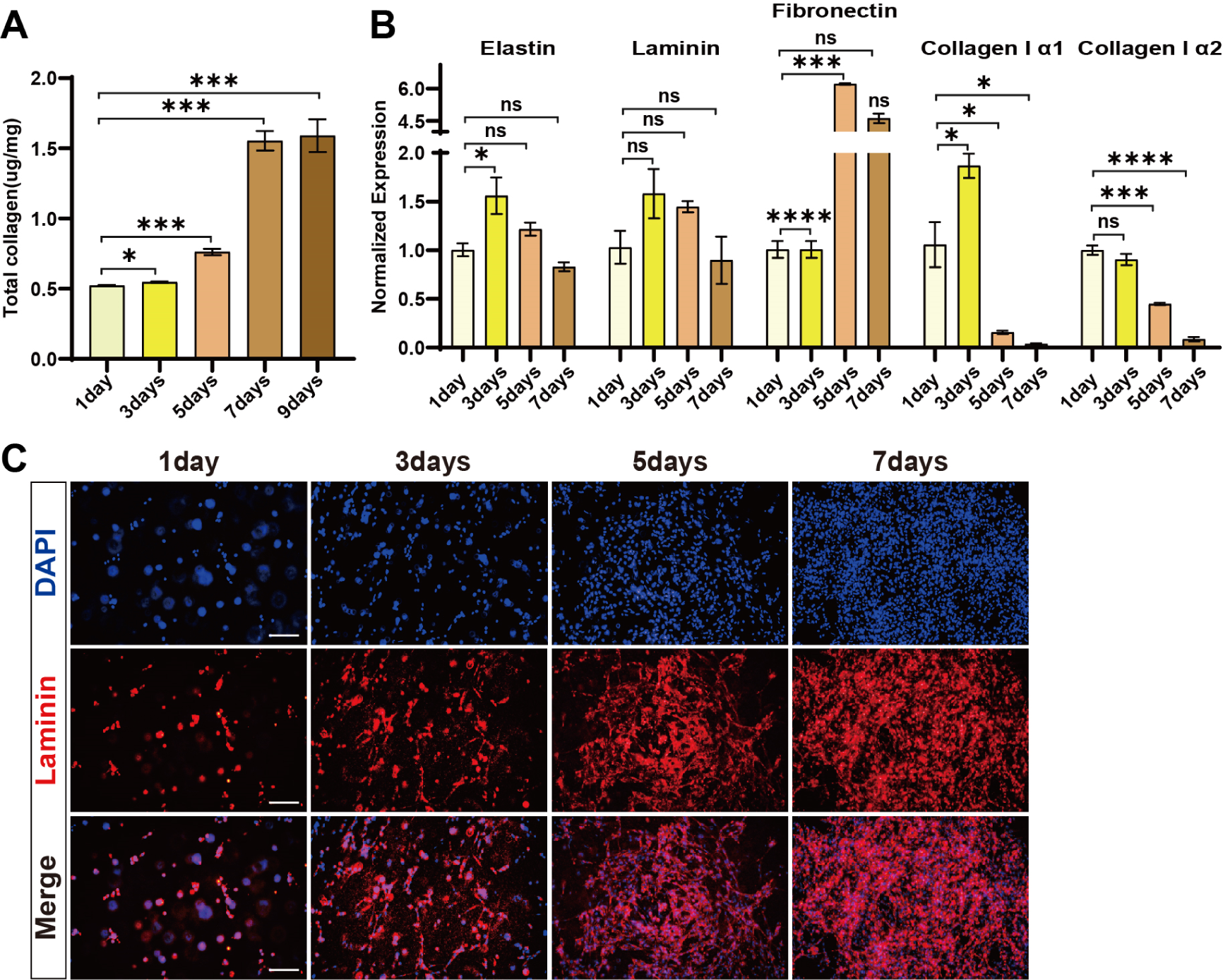
The collagen content and expression of ECM components in cultured meat. (A) Total collagen content of cultured meat at different days of cultivation. Error bars indicate s.e.m, n = 3. *P < 0.05, ***P < 0.001. (B) Expression of ECM-related genes determined by RT-qPCR of cultured meat. Error bars indicate s.e.m, n = 3. *P < 0.05, ***P < 0.001, ****P < 0.0001. (C) Representative Laminin staining of cells in 3D culture on 1day, 3days, 5days and 7days after cell implantation in hydrogel. Scale bar, 100 µm.

### The characterization of molecular changes during myogenic transdifferentiation and fat deposition in cultured meat

To provide insights into the functional shifts during the transdifferentiation from fibroblasts toward muscle, fat, or muscle/fat cells in 3D culture, we further analyzed the transcriptomes from the different populations of cells including the “original fibroblasts” (3D_fibroblast), “myogenic transdifferentiated cells” (3D_MyoD), “adipogenic transdifferentiated cells” (3D+FI), and “myogenic/adipogenic transdifferentiated cells” (3D_MyoD+FI) (Figure 8A). To illustrate the relationship between these cell groups, we conducted unsupervised hierarchical clustering analysis of the entire transcriptome. The findings revealed that the 3D+FI group clustered distinctly from the others, while the 3D_MyoD and 3D_MyoD+FI groups exhibited greater similarity. Moreover, the 3D_fibroblasts formed a distinct sub-cluster on their own (Figure 8B), suggesting that myogenic or adipogenic transdifferentiation drive these cells away from their original fibroblastic state. The Principle Component Analysis (PCA) of the transcriptomes also showed that distinct trajectories of myogenic and adipogenic transdifferentiation routes were derived from the original fibroblasts and finally integrated together into the myogenic/adipogenic cells (3D) (Figure 8C). It indicates that the myogenic and adipogenic signalings could operate simultaneously and separately during the generation of the culture meat composed of muscle and fat.

**Figure 8.**
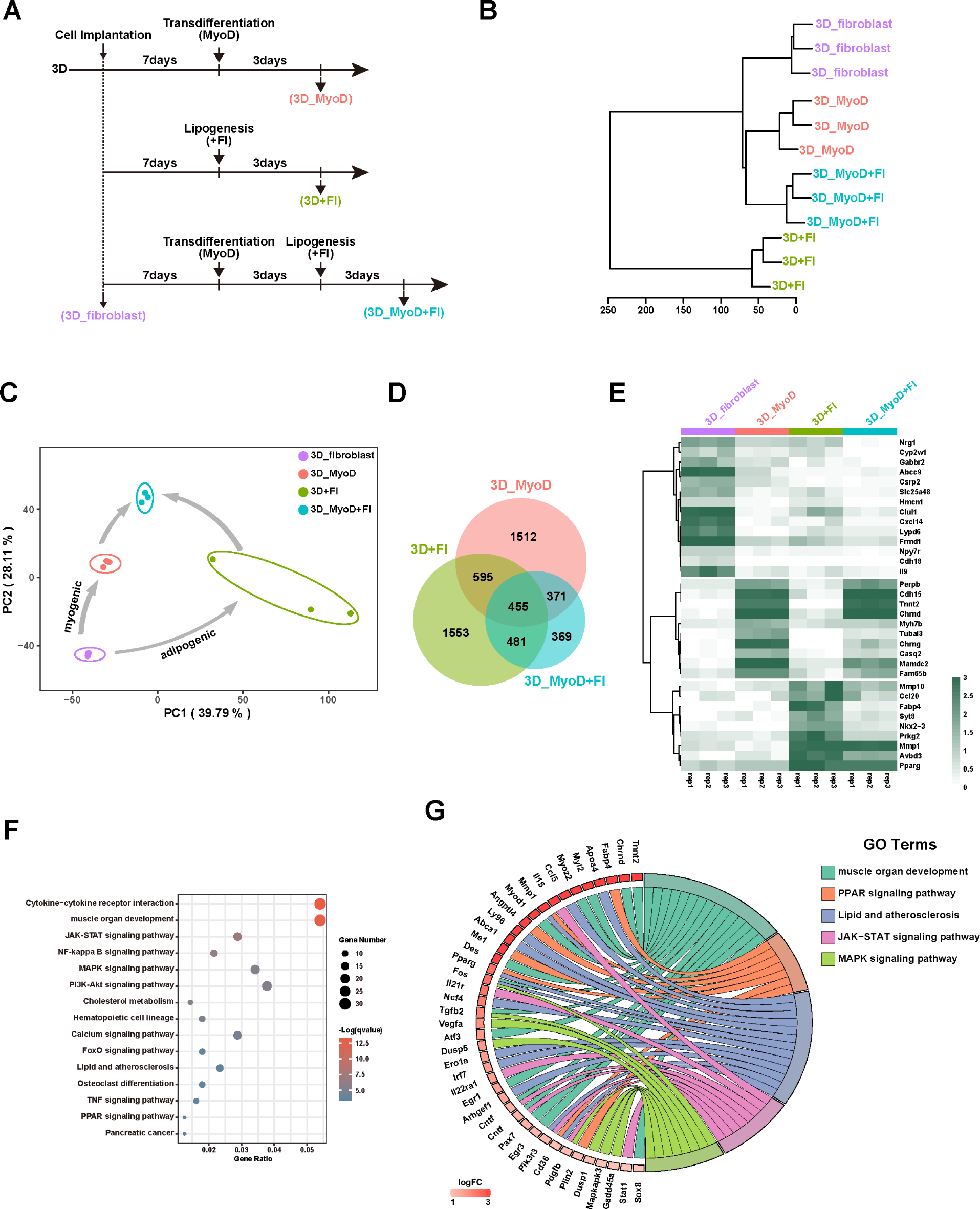
Gene expression profiles during transdifferentiation and fat deposition in 3D. (A) Scheme of the RNA-Seq samples marked by different color. (B) Hierarchical clustering analysis of whole transcriptomes of 3D_fibroblasts, 3D_MyoD, 3D+FI, and 3D_MyoD+FI using Euclidean distance with ward.D cluster method. (C) PCA analysis of transcriptome changes during myogenic transdifferentiation and fat deposition (n= 10,247 genes). The ellipses group include three biological replicates in each cell type. The arrows represent the wiring of gene expression under different conditions. The routes were derived from original fibroblast toward two differentiation routes, namely “myogenic transdifferentiation” and “adipogenic transdifferentiation”. (D) Venn Diagram showing the overlap of differentially expressed genes (DEG) from 3D_MyoD, 3D+FI, and 3D_MyoD+FI compared to the original 3D_fibroblasts. (E) Heat map showed the representative genes differentially expressed between 3D_MyoD+FI and 3D_fibroblast cells (n=3 biologically independent samples). (F) GO analysis of up-regulated DEGs between 3D_MyoD+FI vs. 3D_fibroblast cells. (G) GOChord analysis of the up-regulated genes within representative pathway between 3D_MyoD+FI and 3D_fibroblast cells.

We also compared the differentially expressed genes (DEG) from “3D_MyoD vs 3D_fibroblast”, “3D+FI vs 3D_fibroblast” and “3D_MyoD+FI vs 3D_fibroblast”. The results showed that majority (78%) of DEGs in the 3D_MyoD+FI are overlapped with 3D_MyoD and 3D+FI, indicating that sequential myogenic/adipogenic induction in 3D_MyoD+FI is consistent with myogenic or adipogenic function individually (Figure 8D). The heat map also highlighted the representative myogenic or adipogenic genes that were upregulated in the myogenic, adipogenic, or myogenic/adipogenic cells constitute the culture meat. In contrast, the fibroblast genes were diminished during the transdifferentiation, confirming the loss of fibroblast identity (Figure 8E). In addition, the Gene Ontology (GO) analysis of up-regulated differentially expressed genes (DEGs) in 3D_MyoD+FI cells confirmed that myogenic-specific pathways such as “muscle organ development” and adipogenic-specific pathways such as “PPAR signaling pathway” were enriched. In addition, we also identified several multifunctional signaling pathways such as “JAK-STAT signaling pathway”, “NF-kappa B signaling pathway”, “MAPK signaling pathway” were simultaneously activated during myogenic/adipogenic transdifferentiation, which should have profound effects on both myogenesis(Bakkar and Guttridge, 2010; Jang and Baik, 2013; Keren et al., 2006) and adipogensis (Batista et al., 2012; Bost et al., 2005; Richard and Stephens, 2011) (Figure 8F). The upregulated genes in the representative pathways (such as “*Stat1*” in JAK-STAT signaling pathway, “*Mapkapk3*” in MAPK signaling pathway) were shown in (Figure 8G). In conclusion, the transcriptome analysis of the different types of transdifferentiated cells revealed not only the myogenic- and adipogenic-specific pathways driving the muscle formation and fat deposition respectively, but also several key multifunctional signaling pathways that can promote the cell fate transition and differentiation in different cellular contexts including the muscle and fat tissues.

## Discussion

Mature muscle tissue primarily consists of myofibers (myotubes), which are long, multinucleated cells that contract to generate force and movement. In addition to myofibers, muscle tissue contains a variety of other cell types, including fibroblasts and adipocytes, all of which play important roles in the structure and function of the tissue. The extracellular matrix, which is mainly secreted by fibroblast cells, provides support and structural integrity to the muscle tissue. It is made up of a complex network of proteins and molecules, such as collagen, that provide a scaffold for the cells to attach to and interact with (Franco-Barraza et al., 2016). Together, these components contribute to the unique features of the skeletal muscle tissue and the fresh meat, including its strength, flexibility, and elasticity. It has been and still is challenging to recreate those characteristics in an in vitro condition of cultured meat production. One of the main obstacles in this process is the co-culturing of different cell types with distinct properties. In this study, we overcame this limitation of co-culturing different types of cells by utilizing a single source cell to generate various meat components, including muscle, fat and collagen. Precisely, we employed chicken fibroblasts to produce muscle, deposit fat, and synthesize collagen in a well-controlled and adjustable manner within a 3D setting to produce meat with desirable characteristics.

Fibroblasts are the one of the most common cell types in the animal and could serve as the seed cell for cultured meat production due to their unique and versatile features. First, the fibroblast cells are widely available in the bodies of agricultural animals and could be easily isolated and cultured in vitro. Second, many groups have successfully transformed the chicken fibroblasts into immortalized cell lines (Himly et al., 1998; Pasitka et al., 2023). These cell lines can provide an unlimited cellular resource for cultured meat production and eliminate the need for animal or embryo harvest. Third, fibroblasts are capable of adapting to low serum concentration medium or even serum-free medium, which greatly reduce the culturing cost and risk of serum-bound pathogens (Genbacev et al., 2005; Lohr et al., 2009; Pasitka et al., 2023). Last, the fibroblast cells can also undergo transformation to enable high-density propagation in suspension culture (Bürgin et al., 2020; Fluri et al., 2012; Shittu et al., 2016), which is a crucial step towards scaling up to mass-production. The Food and Drug Administration (FDA) has already approved the use of chicken fibroblast cells for cultured meat production and recently, Eat Just Inc. successfully utilized chicken fibroblasts for the commercial production and sale of cultured meat in Singapore and USA (FDA, 2023). One latest study has also demonstrated the feasibility and consumer acceptance of cultured meat derived solely from native chicken fibroblast cells, indicating a very promising development. However, the resulting meat products do not seem to contain any muscle components (Pasitka et al., 2023). We have previously established an effective strategy for myogenic transdifferentiation, allowing for the production of muscle cells from fibroblast cells of various species in 2D culture (Ren et al., 2022). In the present study, we further enhanced the myogenic transdifferentiation process in 3D, and simultaneously simulated the fat deposition to create cell-based meat that more closely resembles real meat. As a proof-of-concept, we utilized the transgene method to achieve maximum myogenic induction and the final products still retain the foreign transgene fragment in the cells’ genome. It is therefore posing a risk of genetic modified food which is not suitable for mass production. In the next step, other non-transgenic means such as non-integrating vectors, chemical reprogramming, modified RNAs, and recombinant transgene removal techniques will be explored to develop transgene-free end products. Another food safety concern in this study is the use of GelMA hydrogel for culture meat production. Due to its excellent biocompatibility and mechanical flexibility, GelMA-based hydrogel has demonstrated significant potential in scalable 3D cell culture for creating artificial tissue ranging in sizes from millimeters to centimeters. It is widely used in 3D cell culture and tissue engineering for regenerative medicine, but less common in food processing and agricultural applications. Due to its special photo-crosslinking properties, biocompatibility and degradability, it allows this material to be shaped into complex tissue structures by 3D printing or modelling. Many researchers have also used GelMA hydrogel as a scaffold for culture meat production (Jeong et al., 2022; Li et al., 2021; Park et al., 2023). Later research will carefully consider hydrogel as well as other types of scaffold biomaterials for cost-effective and food-safety compliant culture meat production (Bomkamp et al., 2022).

Numerous studies have identified the crucial role of fat in the aroma, juiciness, and tenderness of meat. In general, a very low level of intramuscular fat results in dry meat with plain taste, whereas the high intramuscular fat contents can improve the cooking flavor and greatly increase the value of meat products, such as the high marbling beef from Japanese Wagyu cattle (Gotoh et al., 2018; Motoyama et al., 2016). The fat content of fresh meat mainly come from lipids contained in the fat cells. Thus, closely mimicking the intramuscular fat properties in cultured meat would require co-culturing of muscle cells with fat cells. For example, co-culturing pre-adipocytes with myoblasts may increase the intramuscular fat content, tenderness, and taste intensity of cultured meat (Lau et al., 1996; Pandurangan and Kim, 2015; Zagury et al., 2022). However, co-culture of different cell types is technically challenging, since each cell type grows and differentiates in specific optimized medium. When different cell types are cultured in the same medium, these culture conditions may be sub-optimal for one cell type or the other and result in inefficient cellular growth (Pallaoro et al., 2023). Previous studies have underlined the influence adipocytes growing near muscle cells can impair myogenesis (Seo et al., 2019; Takegahara et al., 2014). Simultaneously or sequentially induction of both myogenic and lipogenic differentiation in the same starting seed cells would resolve these co-culture conflicts and the multi-lineage competent chicken fibroblast cells were chosen to explore this double transdifferentiation strategy. As a proof-of-concept, we successfully transformed chicken fibroblast cells into muscle cells and deposited fat into the same cells in 3D hydrogel matrix. Notably, the intramuscular fat content in the cultured meat could be tailored to any specific value within a certain range. From a nutritional point of view, a direct comparison with the traditional chicken meat was performed. The triglyceride content in the cultured meat is comparable to that of chicken meat and more importantly, the amount of fat could be easily manipulated in order to achieve a more attractive nutritional profile.

In this study, the deposition of fat in the myotubes/myofibers facilitated the storage of significant lipid quantities in transdifferentiated muscle cells, known as intramyocellular lipids. Additionally, we observed Oil Red O staining in the remaining un-transdifferentiated fibroblasts, resembling cells of intramuscular adipocytes (extramyocellular lipids) found within muscle tissue. Hence, current adipogenic induction treatment caused lipogenesis in both the MyoD-transdifferentiated cells and un-transdifferentiated fibroblasts. At present, we don’t know if the fatty acids profile would be different between the two types of origins. The unsaturated fats are conventionally regarded as ‘healthier’ than saturated fats (Berglund et al., 2007; Hamley, 2017). One of the superior assets of cultured meat production is the ability to increase the unsaturated fat levels by adjusting the additive fatty acids, oleic/linoleic acids in this case, in the culture medium. In the future, it requires further comprehensive analysis of the fatty acids profile in the cultured meat to elucidate the fat-associated attributes similar or distinct to real meat.

The current transcriptome analysis during the cellular transdifferentiation also revealed key regulatory signaling pathways control the formation of cultured meat derived from fibroblasts. Notably, we found several multi-functional pathways such as JAK-STAT, MAPK signaling pathways were significantly enriched in the final cells underwent double myogenic/adipogenic transdifferentiation. These signalings could drive the differentiation process of many different types of cells including muscle and adipose tissues (Bost et al., 2005; Jang and Baik, 2013; Keren et al., 2006; Richard and Stephens, 2011), and may be manipulated in current settings to further improve the generation of final cultured meat with tunable fat content.

In conclusion, we have effectively utilized immortalized chicken fibroblasts in conjunction with classical myogenic/adipogenic transdifferentiation approaches within 3D hydrogel to establish a cultured meat model. This model allows for the precise regulation of the synthesis of key components found in conventional meat, including muscle, fat, and ECM. This approach can be readily extrapolated to other species such as pigs and presents promising avenues for the large-scale production of customized and versatile meat products that may cater to varying consumer preferences.

## Materials and Methods

### Cell preparation and inducible myogenic transdifferentiation

The transdifferentiation cells were constructed as described previously (Ren et al., 2022). Briefly, we cloned the chicken full MyoD coding sequence fused in-frame with 3xFlag into a DOX-inducible lentiviral system (Tet-On-MyoD). The wild-type chicken fibroblasts were deal with viral infection and puromycin selection (Invivogen, #ant-pr-1), and finally obtained the transdifferentiation cell lines. In addition, myofibroblasts were isolated from the skin of 10-day-old chicken embryos in the same way as in the previous study (Kosla et al., 2013). Cells were cultured in a 1640 basal medium (Gibco, #C11875500BT) supplemented with 12% fetal bovine serum (FBS) (Gibco, #10270-106) and 1% penicillin-streptomycin (Gibco, #11140050) at 39°C under 5% CO_2_ atmosphere and were given fresh medium every two days. When grown to approximately 80% confluence, the cells were trypsinized and passaged.

### Domestication of cells in low concentration serum medium

The chicken fibroblast cells were domesticated with the progressive concentration of serum. In general, cells were cultured with 12% FBS in a 1640 basal medium and when grown to about 80% confluence, the medium was replaced with 6% FBS medium for further cell culture and passage. The medium was then changed to 3% FBS medium for further cell culture and passage depending on the cell status, and so on. Start again and repeat above steps as soon as the cells grow badly or die. We directly changed the cell culture medium from 12% FBS to 2% CS then reduced it to 1% CS when using chicken serum (CS) (Solarbio, #S9080) medium and the cells could adapt to the medium with low concentration of chicken serum after 3-5 passages.

### Preparation of 3D scaffold and cell culture in 3D matirx

GelMA hydrogels were purchased from Beijing ShangPu for this experiment. Weigh the appropriate amount of GelMA powder into 1640 basal medium, dissolve in a water bath at 70°C for about 30 min, and then filter with 0.22μm sieve. Add 1/8 volume of lithium acylphosphinate salt photoinitiator to the dissolution solution to obtain GelMA hydrogel solution, store at 37°C until usage but no more than 24 h. Fibroblast cells were obtained by trypsin treatment and suspended with GelMA hydrogel solution and gently mixed, followed by treatment of 405nm UV light for 10-20 s to get a 3D hydrogel scaffold. In addition, the hydrogel is secured on a fixation ring for better cell growth and easy movement (Figure S9A and S9B). Place the cell hydrogel complexes in 24-well plate and culture with 1640 medium containing 12%FBS. Transferred to a new well after 12 h, fresh medium was added and changed every 24 h. The cell hydrogel complexes were gently washed three times with PBS buffer and digested by joining collagen II enzyme (Sigma, #10270-106) in incubator at 39°C for 6 min until the cells in the hydrogel package become rounding and shedding. Terminate digest by adding medium and then centrifuged at 1,000 g for 10 min at room temperature (25°C).

### Cell Counting Kit-8 Assay

Cells cultured in 1% CS and 12% FBS were seeded into 96-well plates with 100 μL of medium per well and incubated for 0 h, 24 h, 48 h and 72 h. 10 μL of the cell Counting Kit-8 Assay (CCK-8) solution (Solarbio, #CA1210, China) was then added to each well and incubated for 2 h. The absorbance values at 450 nm were measured by an EnSpire multifunctional spectrophotometer (Perkin Elmer, USA).

### EdU assay

2D cells were cultured in 6-well plates, and 1 mL of the growth medium was added to each well with 0.25 μL of 1.25 mg/mL 5-Ethynyl-2’-deoxyuridine (EdU) (Beyotime, #ST067-1g). After 30 min of incubation, cells were fixed by 4% paraformaldehyde (PFA) for 30 min. Cells were stained with a prepared reaction solution consisting of 1:100 CuSO_4_, 1:4 Tris-HCl, 0.017 g/mL ascorbic acid, 1:1000 Alexa Fluor555 dye and pure water. After 30 min, wash it with PBS, and the nuclei were stained with 50 ng/mL DAPI for 10 min. 3D cultures of cells were cultured in 24-well plate dishes by adding 1mL of the medium with 0.25μL EdU, incubated for 1 h and then fixed with 4% PFA for 24 h. Staining time extended to 1 h and nuclear staining 20 min. Fluorescence images were collected using fluorescence microscopy.

### Cell differentiation

For fibroblast induction into myoblasts, 50ng/ml DOX (Sigma, #D3000000) in 1640 basal medium containing 12% FBS was added for 3 days and the differentiation medium was replaced when the cells reached 80% fusion. In addition, in 3D culture, cells are proliferated for 7 days before changing the differentiation medium when a dense arrangement of cells can be observed under the microscope.

For induction of lipogenic differentiation, the differentiation medium was changed when the fusion rate of cells reached 80% in 2D culture or proliferated for 5 to 7 days in 3D culture. The lipogenesis induction was for 48h with fresh medium changes every 24h. For the differentiation experiments, 1640 with 12% FBS was used as a control, and the lipogenic medium was consistent with a 1:100 fatty acid (’F’ for short) composition of 1:1 oleic acid (Sigma, #O3008, 2 mol oleic acid/mole albumin) and linoleic acids (Sigma, #L9530, 2 mol linoleic acid/mole albumin; 100 mg/mL albumin), and insulin (’I’ for short) (Sigma, #I0516) concentration of 60 μg/mL.

### Immunofluorescence staining

Similar as our previous steps for immunofluorescence staining of cells in 2D culture (Luo et al., 2022), cells were fixed in 4% PFA for 30 min and washed three times with PBS buffer at room temperature, then permeabilizing with 0.5% Triton X-100 on PBS for 10 min and blocking with 10% goat serum at 0.5% TritonX-100 for 1 h. Primary antibodies were diluted with 10% goat serum in 0.5% Triton X-100 at 4°C for 12 h. The primary antibody for MHC (DSHB, #AB2147781) and the primary antibody for Desmin (Sigma, #D8281) and laminin (Sigma, #F1804) were added at a dilution of 1:500 and the primary antibody for Vimentin (DSHB, #AB528504) at a dilution of 1:200. Followed by incubation with Alexa secondary antibody (Invitrogen, #A-21202, #A-21206 and #A-31570) at a 1:500 dilution for 2 h at room temperature. Antibody for α-SMA (Sigma, C6198) was added at a dilution of 1:500 and incubated with 10% goat serum in 0.5% Triton X-100 at room temperature for 2h. Nuclear staining was performed with 50 ng/mL DAPI (Sigma, #D8417) for 10 min. Cells in the 3D culture were fixed for 48 h, the total time of permeabilizing and the blocking was no more than 24 h, and the incubation time of the primary antibody was extended to 24 h, the secondary antibody was 4 h and the nuclear staining 20 min. For MHC detection, fluorescence images were collected using fluorescence microscopy and the MHC^+^DAPI^+^/DAPI^+^ differentiation index was calculated from three or more images using confocal microscopy (Zeiss LSM 800).

### Oil Red O Staining

For Oil Red O staining, cells were fixed with 4% PFA for 30 min or 24 h respectively in 2D and 3D culture. After washing thrice with PBS at room temperature, the cells were soaked in 60% isopropanol for better coloration of Oil Red and then washed for 5 min or 10 min respectively in 2D and 3D culture. The cells were stained with Oil Red O (Sigma, #O0625) for 30 min or 60 min respectively in 2D and 3D. Cells were washed with 60% isopropanol for 30 s or 1 min respectively in 2D and 3D to remove surface staining. Then, cells were washed with distilled water for three times and the stained lipid droplets were visualized using a microscope.

### mRNA extraction and real-time quantitative PCR (qPCR) and library construction

Cells were lysed in Trizol (Simgen, #5301100) and RNA was extracted following manufacture’s recommendations. RNA concentration was measured on NanoDrop2000 (Thermo Scientific, USA). 1 μg of RNA was reverse transcribed using PrimeScriptTM RT reagent Kit (Takara, #RR047A). Real-time quantitative PCR was performed using SYBR Green Mix (Abclonal, #RK21203) following manufacturer’s instructions. Expression was normalized to GAPDH using delta-delta-CT method. For comparisons of the expression, we used a one-tailed Student’s t-test. The error bars indicate the SEM. The RT-qPCR primers are described in Table S1. The RNA-Seq library-preparation protocol was based on the NEBNext Ultra™ RNA Library Prep Kit for Illumina (NEB, #E7530L). Insert size was assessed using the Agilent Bioanalyzer 2100 system and qualified insert size was accurate quantification using StepOnePlus™ Real-Time PCR System (Library valid concentration>10nM) and then paired-end sequencing using an Illumina platform. RNA-Seq was performed on triplicates for each sample.

### RNA-Seq data analysis

The RNA-Seq raw data were firstly trimmed adapters by trim_galore software (Bolger et al., 2014) and then the clean data of RNA-Seq was mapped to chicken genome (Ensembl, GRCg7b) using hisat2 (Kim et al., 2019) with default parameters. Because of using paired-end reads, the concordant unique mapping reads/pairs were kept based on the mapping flags. The duplications were removed based on the coordinates of the reads/pairs. The de-duplication unique mapping reads/pairs were used for further analysis in this study. Read counts for each sample were computed with the featureCounts (Liao et al., 2014) software and the RefSeq gene annotation for chicken genome assembly is GRCg7b. The transcript per million (TPM) were normalized using reads counts. Differential expressed genes (DEGs) of RNA-Seq data were analyzed using DESeq2 called by q<0.01 and fold change >2 thresholds. The chicken genes were transformed to homologue human genes using ensemble bioMart database and the gene ontology in this study was conducted in Metascape and the enriched top pathways were shown. All plots were generated by R (v4.0.3).

### Measurement of total collagen content in cultured meat

Total collagen content was determined by the concentration of hydroxyproline. The medium was replaced with fresh medium in the incubator for 2 h before the assay, washed three times with PBS. According to the instructions of the Hydroxyproline Assay Kit (Jiancheng, #A030-1-1, China), one volume of saline was added in dried cultured meat at a 1:1 ratio of weight and volume. After mechanical homogenization in ice-water bath, the digestion solution was incubated at 37°C in a water bath for 4 h. Hydroxyproline content was measured by the absorbance at 550 nm using an EnSpire multifunctional spectrophotometer (Perkin Elmer, USA). For each measurement of cultured meat, the hydroxyproline content of the corresponding blank cell-free hydrogel was substracted. The amount of collagen was calculated from the hydroxyproline concentration with a conversion factor of 7.25 in μg/mg wet tissue (Vasanthi et al., 2007; Zheng et al., 2021).

### Measurement of triglyceride content in cultured meat

The triglyceride content in cultured meat was measured using the kit (Solarbio, #BC0620, China). Before the assay, the culture system is washed three times with PBS and uniformly added to the same medium containing 12% FBS in the incubator for 2 h, after which it is removed and washed three more times with PBS, the cultured meat is taken out and churned, and the precipitate is obtained by centrifugation, which should be well air-dried. The precipitate was weighed and tirzol was added to lyse well for 2 h. A mixture of n-heptane and isopropanol in a ratio of 1:1 was then added and shaken and mixed. Then added potassium hydroxide and full shaken to produce glycerol and fatty acids, then the other reagents were added in sequence according to the instructions. The final triglyceride content was measured the specific light intensity at 420 nm, as described previously.

### Emission scanning electron microscope

The cell hydrogels were washed three times with PBS and then fixed in 4% PFA for 1 h. Samples were randomly clamped into spiking trays, rapidly frozen in liquid nitrogen and then images were randomly collected using the emission scanning electron microscope (HITACHI SU8010, Japan).

### Statistical analyses

Statistical analyses were performed using the Graph-Pad Prism software. For normally distributed data sets with equal variances, a two-sample t-test was used. The significance of differences was provided in figure legends.

## Acknowledgements

This work was supported by National Natural Science Foundation of China (31771617), Taishan Scholar Program, Scientific Research Innovation Team of Young Scholar of Shandong.

