## Supplementary file for "Transdifferentiation of fibroblasts into muscle cells to constitute cultured meat with tunable intramuscular fat deposition"

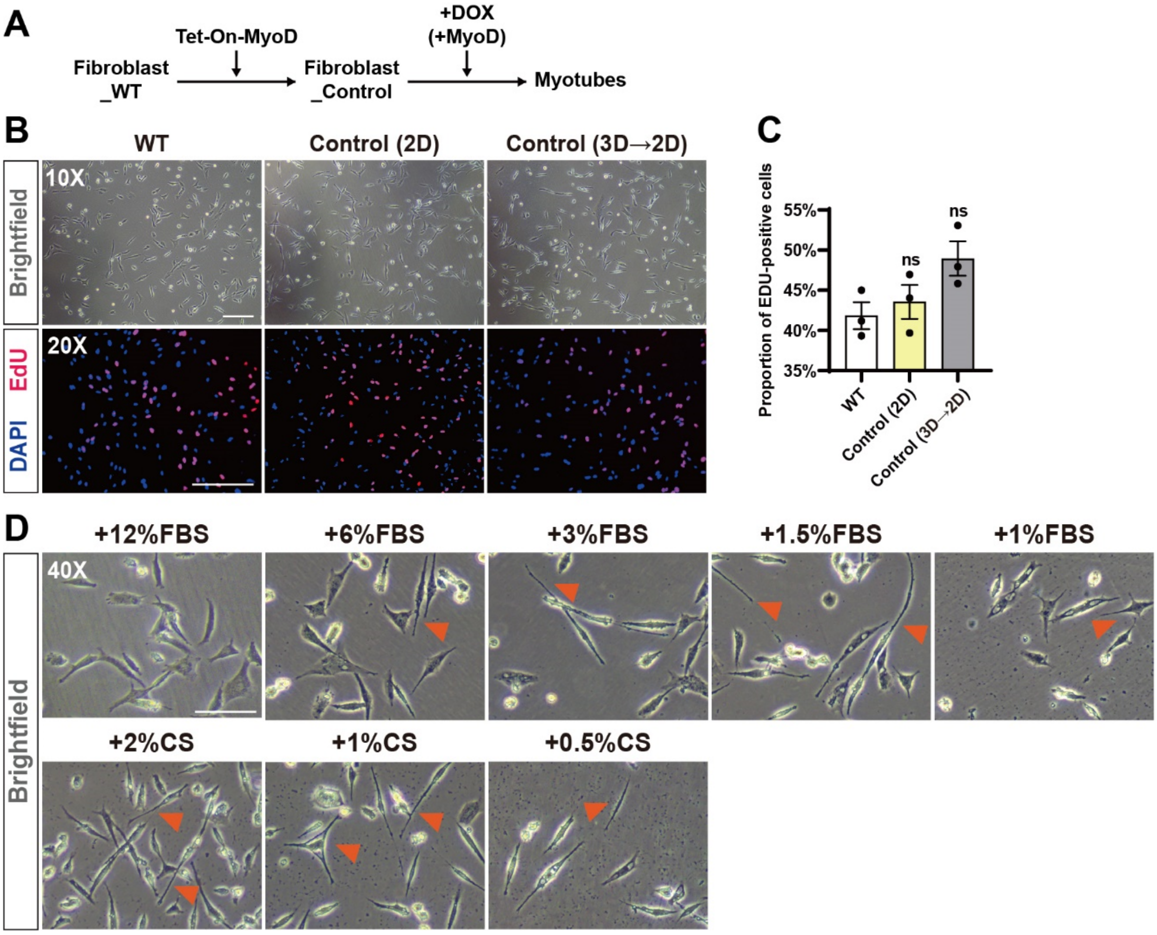


**Supplementary Figure 1.** (A) Scheme of the MyoD-induced transdifferentiation. (B) Morphology and EdU staining of chicken fibroblasts under different conditions. Scale bar, 200 μm. (C) Quantification of the proportion of EdU-positive cells in Figure B. Error bars indicate s.e.m, n = 3. (D) Cellular morphology of chicken fibroblasts under different low-serum conditions. Orange triangles mark the sharper and smoother morphology of cell edge contours. Please note that the cells showed abnormal morphology in the lowest serums of 1% FBS and 0.5% CS. FBS: Fetal bovine serum. CS: Chicken serum. Scale bar, 100 μm.

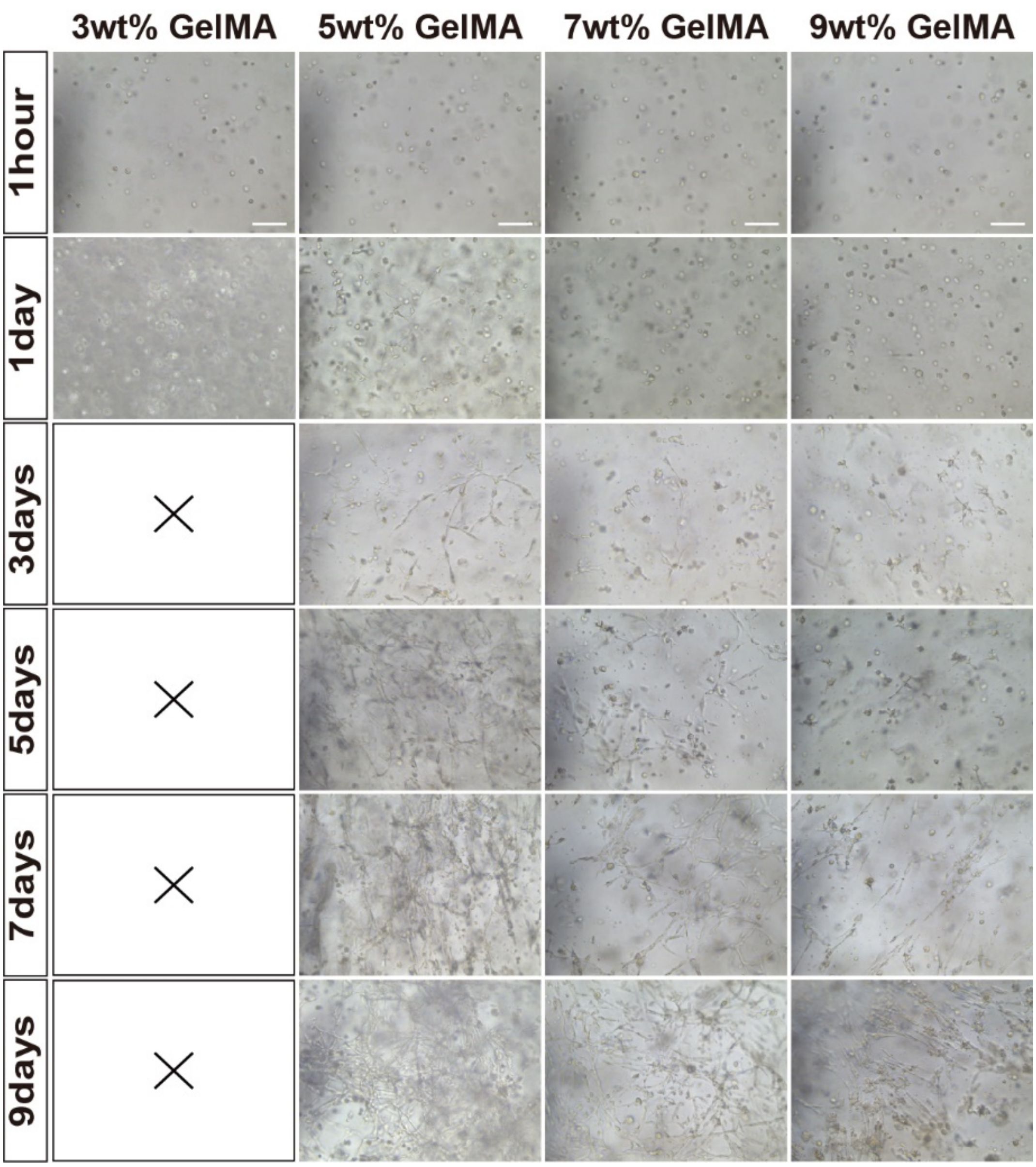


**Supplementary Figure 2.** Morphological changes of cells implanted and cultured on four different concentrations of hydrogels at 3wt%, 5wt%, 7wt% and 9wt% for different times, and the 3wt% hydrogel collapsed after the second day of growth. Scale bar, 100 μm.

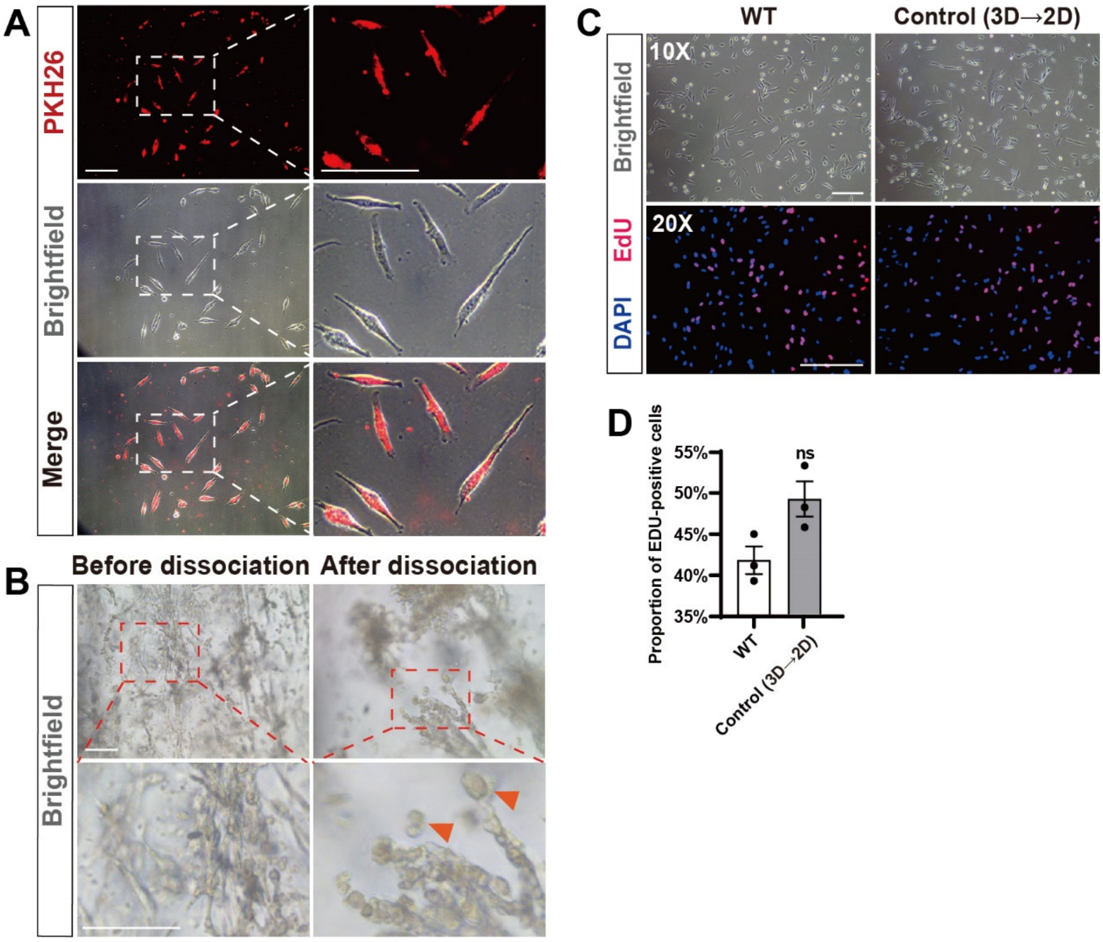


**Supplementary Figure 3.** (A) Brightfield and red fluorescent images of chicken fibroblasts in 2D culture after PKH26 labeling. Scale bar, 100 µm. (B) Morphology of cells in GelMA hydrogels cultured in 3D before and after dissociation. Scale bar, 100 µm. (C) Morphology and EdU staining of chicken fibroblasts under different conditions. 3D→2D indicates that cells isolated from 3D (Figure B) were re-cultured in 2D. Scale bar, 200 µm. (D) Quantification of the proportion of EdU-positive cells in Figure C. Error bars indicate s.e.m, n = 3.

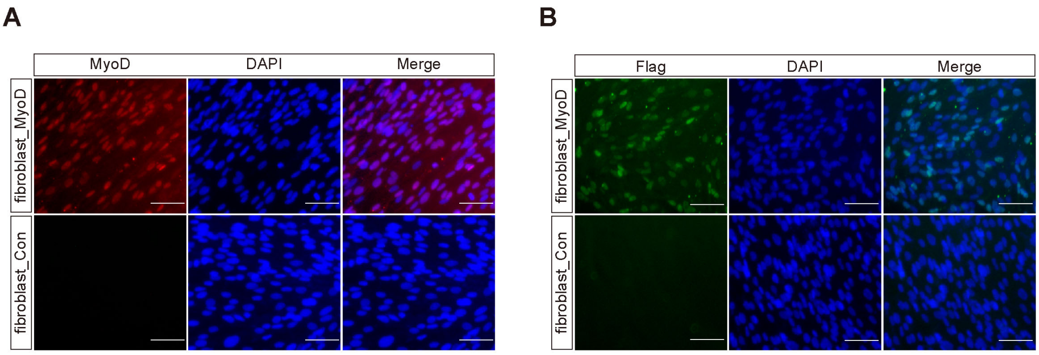


**Supplementary Figure 4.** Expression of MyoD upon Dox treatment. The MyoD-3xFlag was fused in frame and under the control of Tet-On system. **(**A) Representative immunofluorescence staining of MyoD in fibroblast_MyoD and fibroblast_Con. Scale bar: 50μm. Scale bar, 50 μm. (B) Representative immunofluorescence staining of Flag in in fibroblast_MyoD and fibroblast_Con. The anti-Flag immunostaining indicate the exogenous MyoD transgene expression. Scale bar, 50 μm.

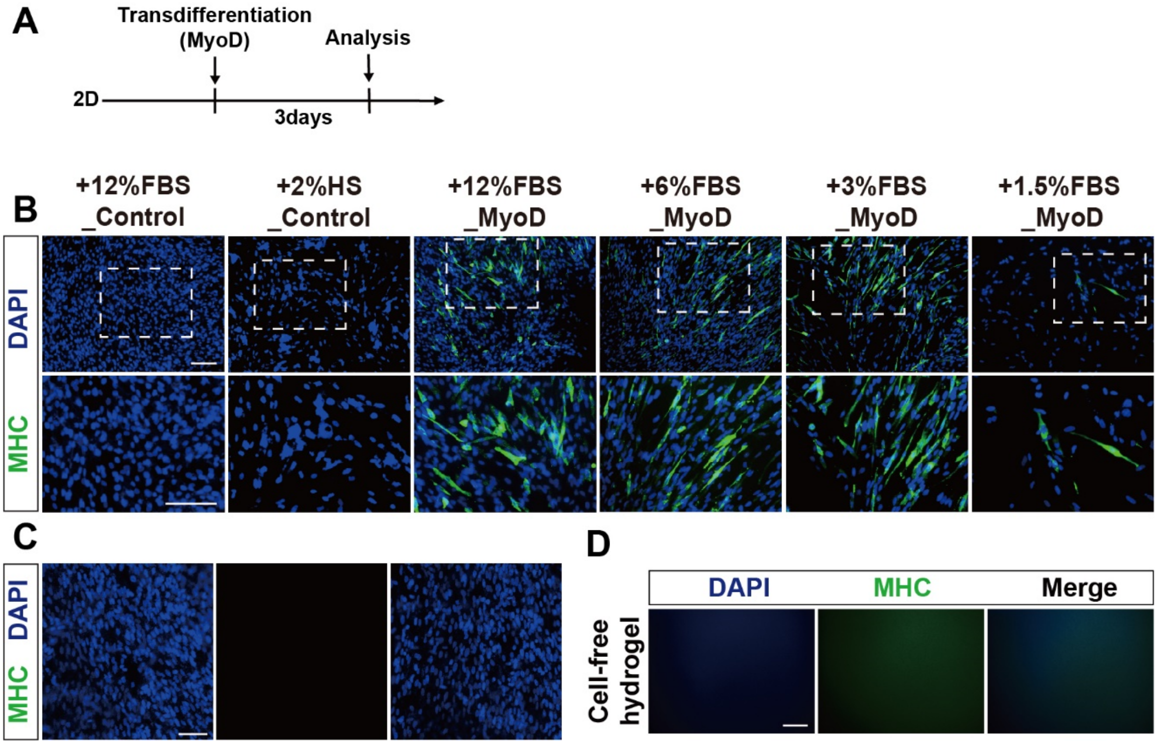


**Supplementary Figure 5.** (A) Experimental design for fibroblast myogenic transdifferentiation in 2D culture. (B) MHC staining demonstrates the myogenic capacity of chicken fibroblasts. Scale bar, 100 µm. (C) Immunofluorescence staining of MHC in 3D cultured chicken fibroblasts without activation of MyoD factor as a negative control. Scale bar, 100 µm. (D) Immunofluorescence staining MHC in cell-free GelMA hydrogels as the control. Scale bar, 200 µm.

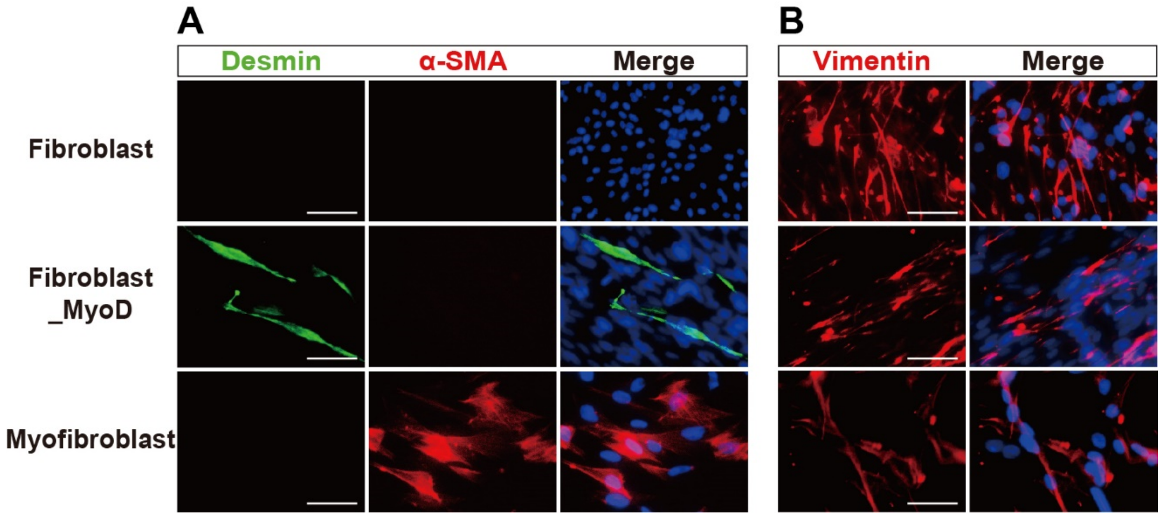


**Supplementary figure 6.** (A) Immunofluorescence staining of 2D cultured cells showed that the skeletal muscle marker Desmin was expressed only in the transdifferentiated cells but not in fibroblasts or myofibroblasts, and the myofibroblast marker α-SMA was expressed only in the myofibroblasts, but not in fibroblasts or transdifferentiated cells. Scale bar, 50 μm. (B) Immunofluorescence staining of 2D cultured cells showed that the fibroblast marker Vimentin was abundantly expressed in fibroblasts but greatly reduced in MyoD transdifferentiated cells. Scale bar, 50 μm.

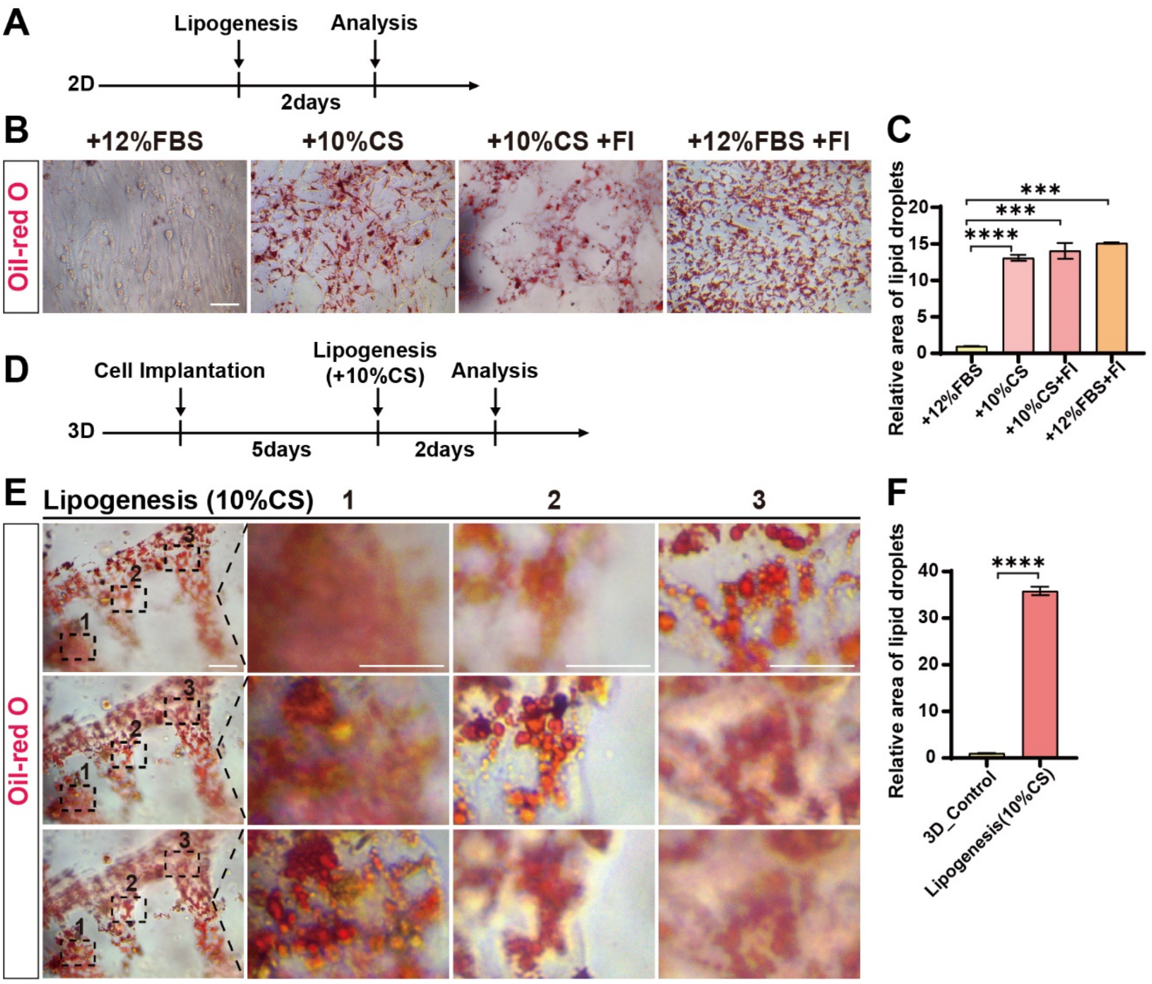


**Supplementary Figure 7.** (A) Experimental design of fibroblast lipogenic differentiation in 3D culture. (B) Oil-Red O staining of lipids in 2D under different conditions. Scale bar, 100 µm. (C) Relative area of lipid droplets in figure C. Error bars indicate s.e.m, n = 3. ***P < 0.001, ****P < 0.0001. (D) Experimental design for fibroblast lipogenic differentiation in 3D culture induced by chicken serum (CS). (E) Oil-Red O staining of 3D culture of cells after lipogenic induction and representative images were taken consecutively at different focal planes in the same position. "1", "2", "3" are magnifications of the corresponding areas. Scale bar, 100 µm. (F) Relative area of lipid droplets in figure E. Error bars indicate s.e.m, n = 3. ****P < 0.0001.

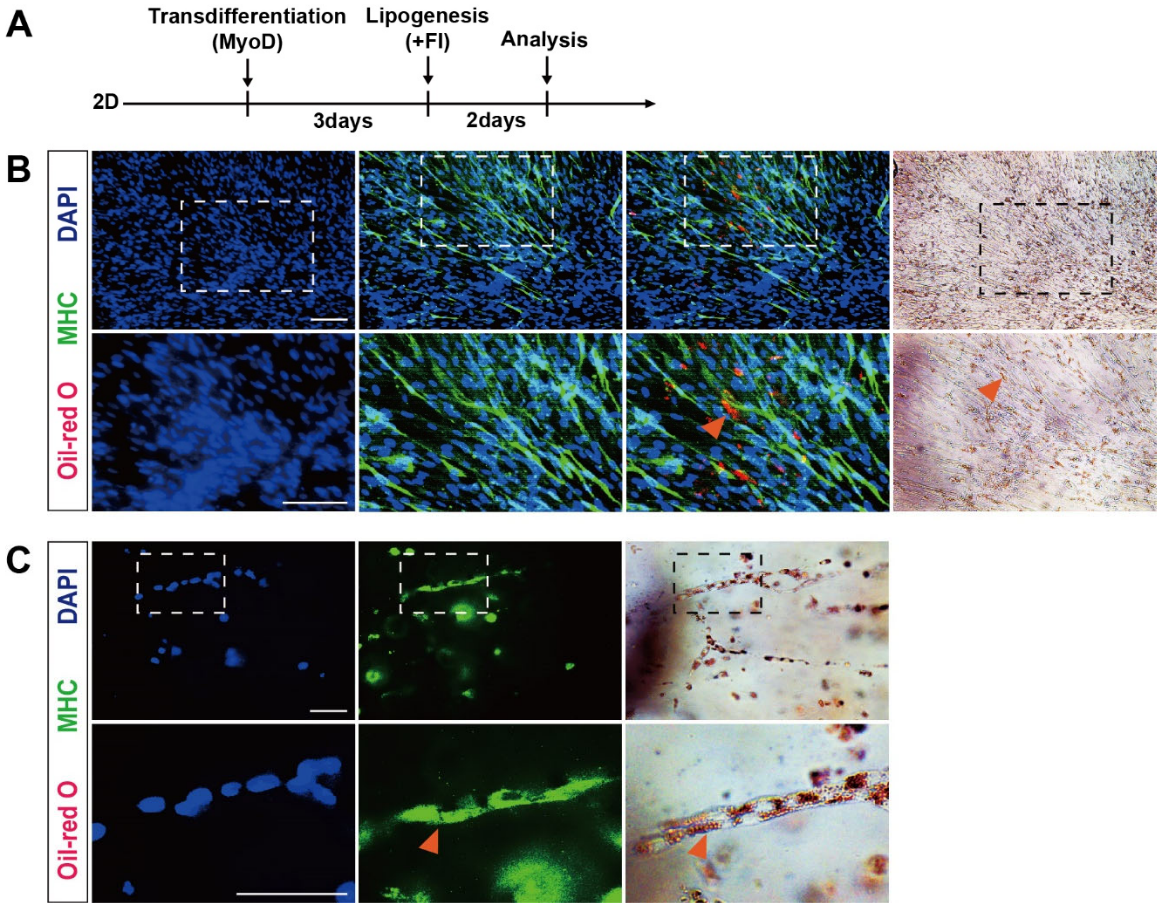


**Supplementary Figure 8.** (A) Experimental process for sequential myogenic/lipogenic stimulation in 2D culture. (B) MHC staining and Oil Red O staining of cells with 2D induction of myogenesis/lipogenesis, with triangular arrow indicating that Oil Red O-labelled lipid droplets are shown under the red fluorescent channel. Scale bar, 100 μm. (C) Immunofluorescence staining and Oil Red O staining demonstrating lipid deposition inside the transdifferentiated muscle cells, with triangular arrow marks the location of the lipids within the muscle cell. Scale bar, 100 μm.

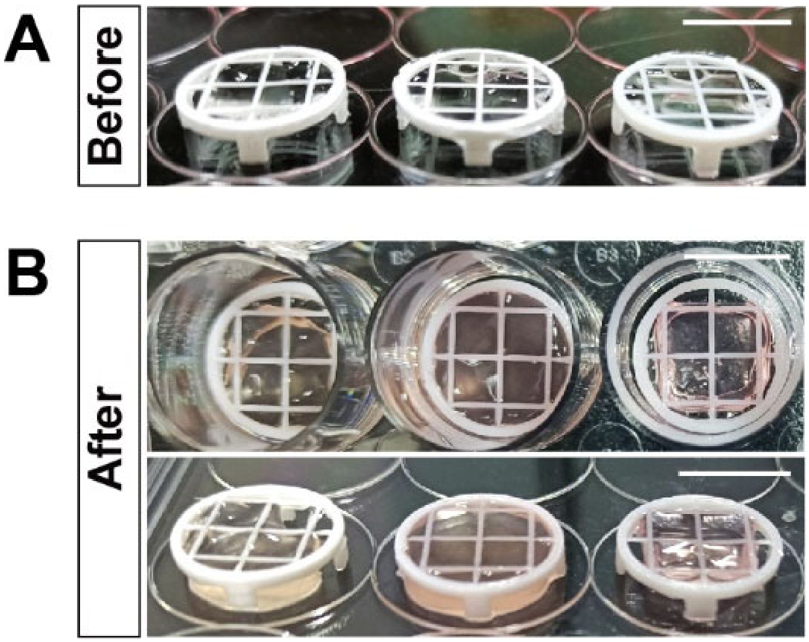


**Supplementary Figure 9.** Cell-GelMA 3D culture units and macroscopic morphology after 7 days of culture, with white plastic frame as the fixation ring. Scale bar, 1 cm.

**Supplementary table 1. List of primers of qPCR**

| Gene Name | Forward primer | Reverse primer |
|---|---|---|
| *Gapdh* | *TCGGAGTCAACGGATTTGGC* | *ATAGTGATGGCGTGCCCATT* |
| *MyoD* | *ACTACAGCGGGGAGTCAGAT* | *GCTTCAGCTGGAGGCAGTAT* |
| *MyoG* | *AGCCTTCGAGGCTCTGAAAC* | *AAACTCCAGCTGGGTGCTC* |
| *Myh15* | *AGATAAAGGAACTACAGGCTCGT* | *CGCCAGCTTCAGGAACTCA* |
| *CKM* | *ACCTGGACCCCAAATACGTG* | *TCGAACAGGAAGTGGTCGTC* |
| *Desmin* | *GGAGATCGCCTTCCTCAAGA* | *CAGGTCGGACACCTTGGATT* |
| *Six1* | *ACTGCTTCAAGGAGAAGTCG* | *TTCTCCGTGTTCTCCCTCTC* |
| *Thy-1* | *TGTCATCCTGACAGTGCTGC* | *GGTAGAGGCACACCAGGTTC* |
| *TGFβ-1* | *GAGCTGTACCAGGGTTACG* | *GAAGCCTTCGATGGAGATG* |
| *TGFβ-3* | *CTCCCCGAGCACAATGAGT* | *TATATGCTCATCTGGCCGCA* |
| *Smad3* | *GCAAGATCCCACCAGGATG* | *GAGGTGCAGCTCAATCCAG* |
| *Pparg* | *TGCCAAGCATTTGTAT* | *TGCGAATTGCTACTTCTTTGTT* |
| *Znf423* | *CCAGTGCCCACAGAAGTTCT* | *CCACTGTGCCACCATCAAGT* |
| *Fabp4* | *CAAGCTGGGTGAAGAGTTTGATG* | *TCGTAAACTCTTTTGCTGGTAAC* |
| *Gpd1* | *GGCTTTTGCCAAGACTGGGAA* | *GGTTTGCCCTCATAGCAGATCTG* |
| *Collagen I α1* | *GTCCTGCTGGATTTGCTGG* | *GAAACCAGTAGCACCAGGG* |
| *Collagen I α2* | *TGATCCATCTAAAGCGGCTG* | *TTTGCCAGGGTGACCATCTT* |
| *Laminin* | *CGCGATTTCTGATTTTGCCG* | *CATTGCAGTCACAAGGCAAG* |
| *Fibronectin* | *GTGCTACGACGATGGGAAAA* | *GCAGTTGACGTTGGTGTTTG* |
| *Elastin* | *CTACTGGGACAGGTGTTGGA* | *CACCATAGGCTCCTGCCTT* |

**Supplementary videos**

Supplementary video 1. MHC+ staining of cultured meat (myogenic).
Supplementary video 2. MHC+ staining of cultured meat (myogenic) (different view).
Supplementary video 3. Oil-Red O staining of cultured meat (lipogenic).
